# Astrocytic PIEZO activation by heartbeat sound promotes extracellular matrix and maturation in human brain organoids

**DOI:** 10.1101/2025.10.21.683816

**Authors:** Alexandra Ferreira Oliveira, Florian Rambow, Samira Makhzami, Yury A. Nikolaev, Angelo Novak, Olga Shevchuk, Hannah Voss, Daniel R. Engel, Paul A. Schult, Matthias Pillath-Eilers, Britta Kaltwasser, Ulf Brockmeier, Christoph Kleinschnitz, Dirk M. Hermann, Egor Dzyubenko

## Abstract

Heartbeat sound is one of the first rhythmic stimuli encountered by the developing human brain, yet its biological role has remained unexplored. Here we show that heartbeat sound promotes structural and functional maturation of human cortical organoids with a physiological astrocyte-to-neuron ratio. Continuous stimulation with heartbeat expanded the extracellular space (ECS), visualized by super-resolution 3D STED and 2-photon shadow imaging, and increased extracellular matrix (ECM) synthesis, detected by LC-MS/MS proteomics. Heartbeat sound promoted neuronal and astrocytic differentiation, synaptogenesis, and organoid maturation, as shown by single-cell RNA sequencing, synaptic marker immunohistochemistry, and high-density multielectrode electrophysiology. Transcriptomic analyses demonstrated astrocyte-specific expression of mechanosensitive PIEZO channels. Using Ca^2+^ imaging, we confirmed that low-frequency sound stimulation, including heartbeat, activates PIEZO-mediated Ca^2+^ responses, which in turn upregulate astrocytic ECM synthesis. Experiments with biomimetic hydrogels demonstrated that physiological matrix stiffness attenuates PIEZO activation, suggesting a mechanoprotective mechanism. Our findings demonstrate a fundamental mechanism for inducing ECM synthesis and neural differentiation through astrocytic PIEZO activation by heartbeat sound, suggesting that interoception of the internal acoustic environment is an underappreciated driver of human brain development.

**Highlights:**

- Heartbeat sound activates PIEZO ion channels
- Astrocytic PIEZO activation promotes extracellular matrix synthesis
- Heartbeat sound stimulates neural differentiation and brain organoid maturation

**Graphical Abstract:** 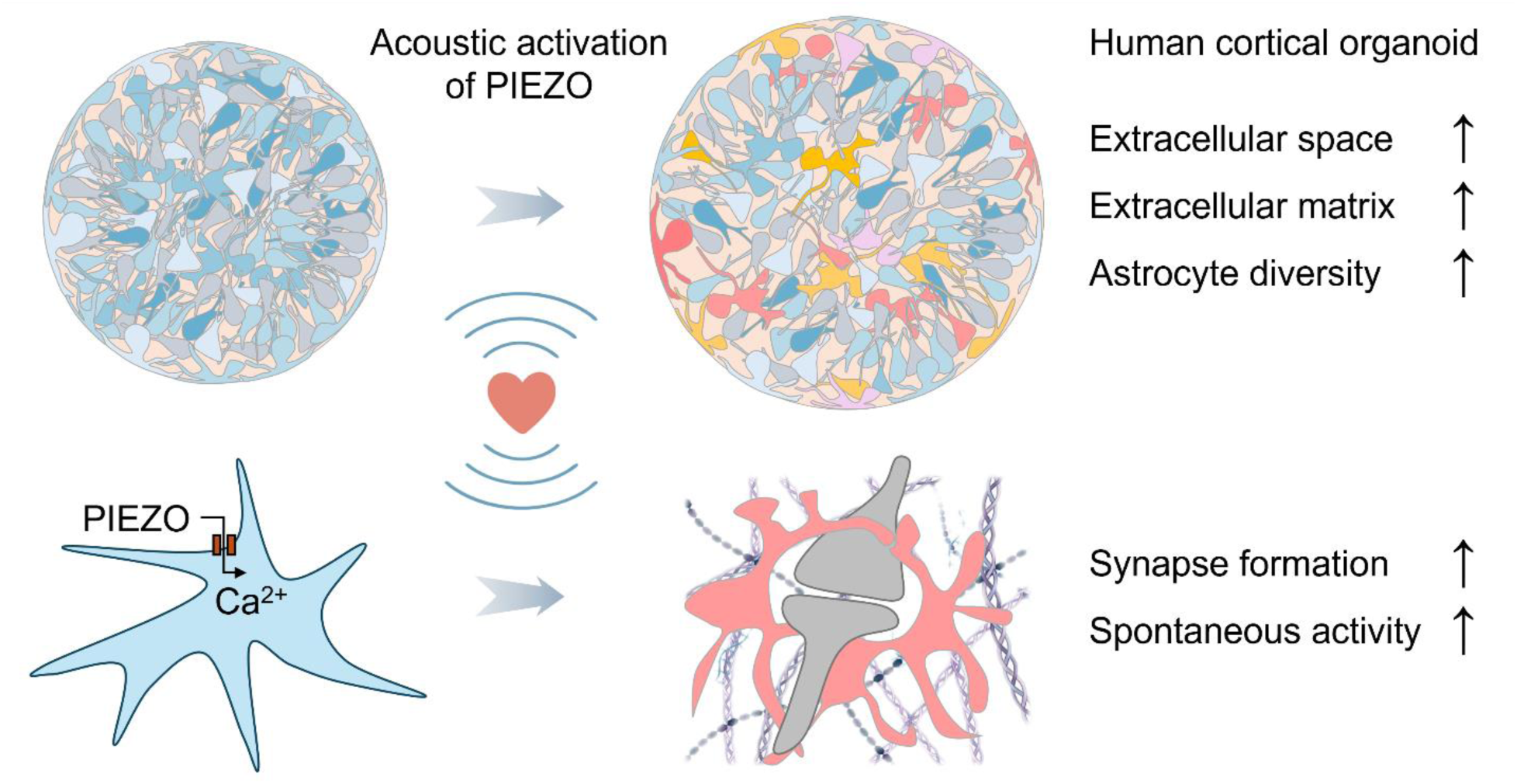

## Introduction

Mother’s heartbeat is one of the first sounds that a human fetus hears starting the third trimester of pregnancy (Querleu et al., 1988). Yet, even before the auditory system fully develops, the embryo is exposed to acoustic stimulation, which is dominated by sound frequencies below 200 Hz including those elicited by maternal heartbeat (Dealessandri and Vivalda, 2018; Vince et al., 1982). While the high frequencies are strongly attenuated inside the womb, low frequency sounds can be as loud as 100 dB (Dealessandri and Vivalda, 2018; Gerhardt et al., 1990). Therefore, the acoustic environment of an embryo is defined by low-frequency sound waves generating persistent mechanical oscillations that rhythmically stimulate developing tissues.

In mammalian tissues, mechanical oscillations are mainly detected by PIEZO ion channels (Coste et al., 2010; Zhao et al., 2018), which have been implicated in interoceptive regulation of breathing (Tort et al., 2025) and blood pressure (Jammal Salameh et al., 2024; Zeng et al., 2018). PIEZO1 and PIEZO2 are mechanically activated ion channels that share a common activation mechanism gated by membrane tension. While PIEZO2 is predominantly expressed by somatosensory neurons, PIEZO1 is also found in glial (Chi et al., 2022), vascular (Ranade et al., 2014), and immune cells (Solis et al., 2019).

The molecular mechanisms of sound-driven mechanotransduction remain underinvestigated. Although the possibility of inducing cellular responses by sound stimulation has been demonstrated (Kumeta et al., 2025; Xu et al., 2024), the physiological relevance and mechanistic understanding of such stimulation has remained elusive. We hypothesized that heartbeat sound provides a natural mechanical stimulus that shapes embryonic brain development.

For experimental studies of human neurodevelopment, brain organoids derived from human induced pluripotent stem cells (hiPSCs) have become the model of choice. Originally designed to recapitulate cortical organization (Lancaster et al., 2013) and to model developmental disorders, organoid systems have since been adapted to study interactions between brain regions and to investigate connectivity-related diseases such as epilepsy (Onesto et al., 2024). Despite these advances, current organoid models do not capture the acoustic environment during early brain development. In this study, we investigated how mechanostimulation with heartbeat sound shapes cellular physiology and functional maturation in human cortical organoids, focusing on the brain extracellular matrix (ECM), neural cell differentiation, synaptogenesis, and neuronal activity.

## Results

### Heartbeat sound expands the extracellular space in astrocyte-enriched organoids

Astrocyte-enriched cortical organoids were generated by co-culturing neural precursor cells (NPCs) with induced astrocytes (iAstrocytes) using a modified growth factor-based protocol (Paşca et al., 2015) (**Fig. 1A**). By day 45, they developed ramified astrocytes, glutamatergic synapses, and showed increased ECM expression compared with non-enriched cortical organoids (**Fig. S1**). Cortical organoids with and without astrocyte enrichment were exposed to a human heartbeat audio recording for 31 days of maturation, with ambient noise as control. Hydrophone measurements in cultivation medium (**Fig. 1B**) were used to tune the applied heartbeat sound to physiological acoustic environment of an embryo during gestation (**Table 1**).

**Figure 1.**
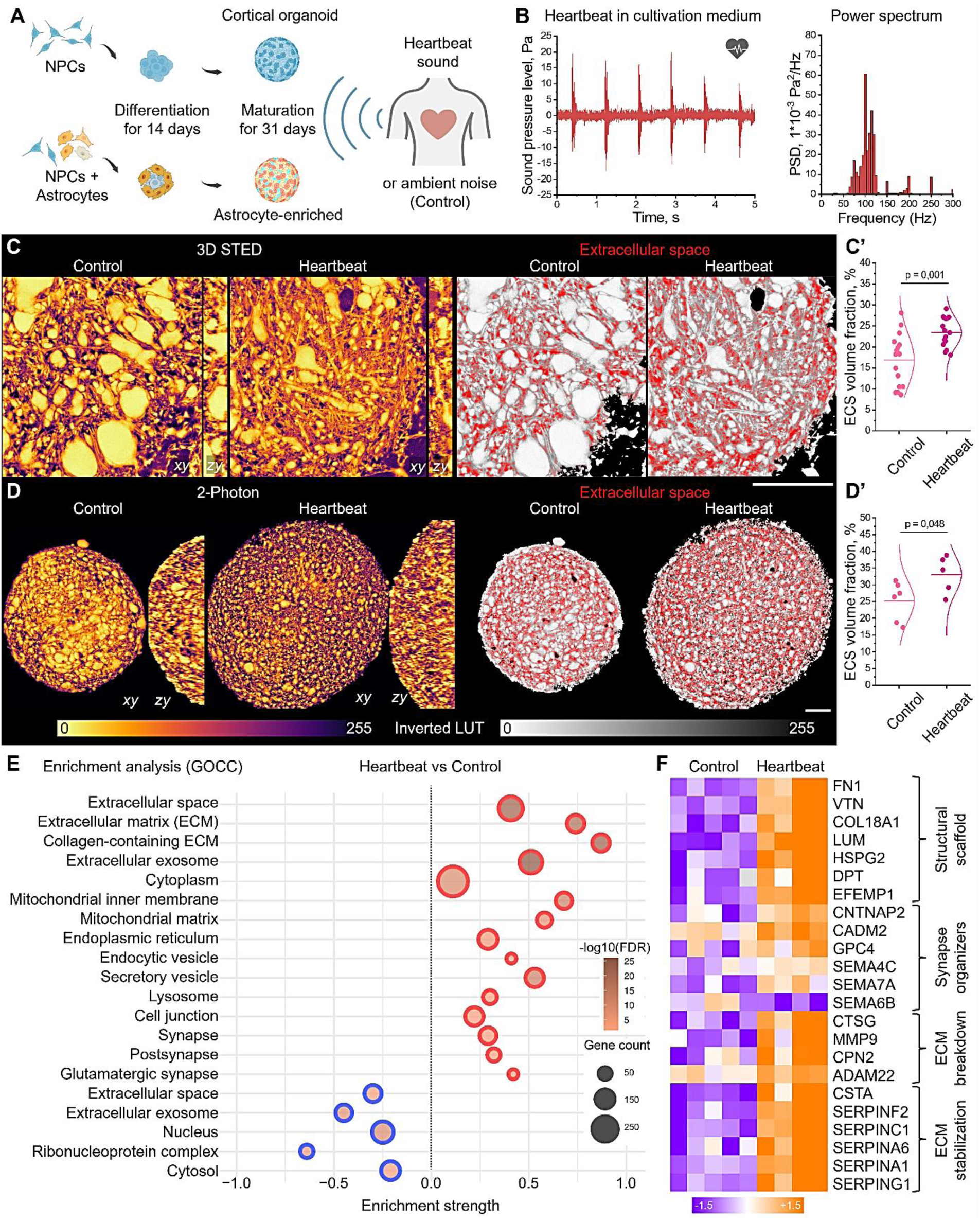
Stimulation with heartbeat sound increases ECS volume and ECM expression in astrocyte-enriched human cortical organoids. (**A**) Generation of organoids and exposure to heartbeat. (**B**) Hydrophone recording and power spectrum of the applied heartbeat sound in cultivation medium. (**C**) Superresolution 3D STED shadow imaging and ECS segmentation (red) in live organoids. (**C’**) ECS volume fraction comparison, t-test (n=10-15). (**D**) 2P shadow imaging and ECS segmentation (red) in live organoids. (**D’**) ECS volume fraction comparison, t-test (n=5-6). (**C**, **D**) Images are xy and zy projections; LUT inverted (see **<u>Fig S2</u>** for details). Scale bars, 50 µm. (**E**) Gene ontology cellular component (GOCC) enrichment analysis of protein expression in heartbeat-stimulated vs control astrocyte-enriched organoids, based on significantly regulated proteins (p < 0.05, fold change difference > 1.5, t-test), measured by LC-MS/MS. (**F**) Heatmap shows normalized, centered (mean subtraction) log2 protein abundances of selected ECM proteins regulated by heartbeat sound. Other regulated ECM proteins are shown in **<u>Fig S4</u>.**

**Table 1.**
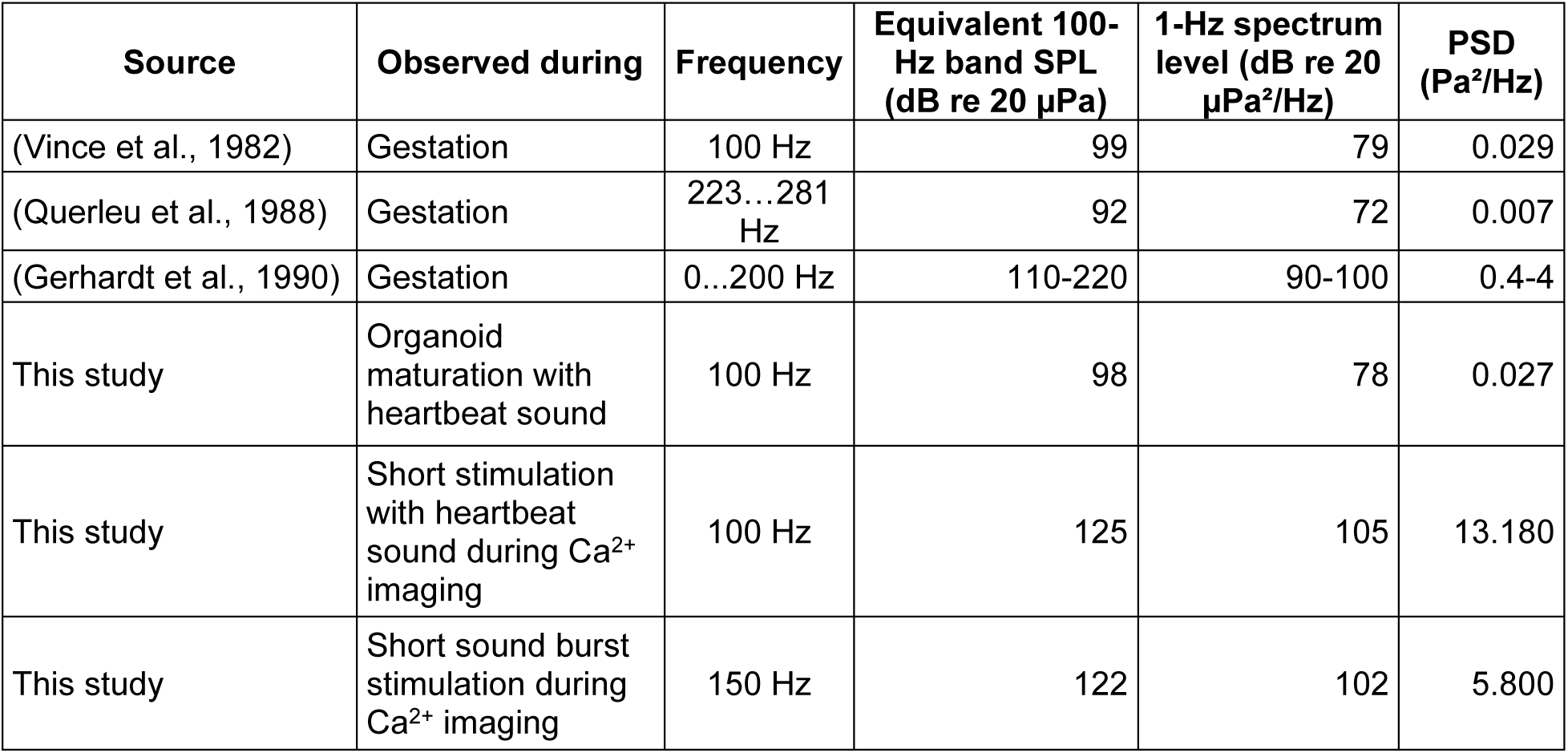
Uterine sounds and acoustic stimulation of organoids.

Heartbeat sound increased the extracellular space (ECS) volume fraction in astrocyte-enriched organoids by 39% (23.4±3.5 vs 16.8±6.2 %, mean±SD) (**Fig. 1C, C’**), shown by 3D STED superresolution shadow imaging (SuSHi) (Tonnesen et al., 2018), explained in (**Fig. S2**). This observation was confirmed with 2-photon (2P) shadow imaging indicating a 31% (33.1±5.6 vs 25.2±5.8 %, mean±SD) ECS volume increase in astrocyte-enriched organoids stimulated with heartbeat sound (**Fig. 1D, D’**). Higher ECS volume fractions measured with 2P are due to lower resolution compared with 3D STED (**Table S1**) and consequent under-detection of small cellular structures. No major changes in the ECS volume were observed in organoids without astrocyte enrichment (**Fig. S3**).

### Heartbeat sound promotes ECM synthesis in astrocyte-enriched organoids

Because the expanded ECS needs to be stabilized by an extracellular scaffold to maintain tissue integrity and diffusion (Tønnesen et al., 2023), we next analyzed protein expression using LC-MS/MS. Gene Ontology Cellular Component (GOCC) enrichment analysis comparing heartbeat-stimulated and control organoids revealed ECM among the top regulated terms (**Fig. 1E)**. In astrocyte-enriched organoids, heartbeat sound stimulation upregulated multiple ECM scaffolding proteins, among them fibronectin (FN1) and vitronectin (VTN), and synapse organizing molecules including glypican 4 (GPC4), neuroligins (NLGN1, NLGN2), and plexin-B (PLXNB1) (**Fig. 1F** and **Fig. S4).** In organoids without astrocyte enrichment, we found no major changes in ECM expression following heartbeat sound stimulation (**Fig. S4**).

### Heartbeat sound promotes neural differentiation in astrocyte-enriched organoids

The brain ECM is jointly produced by neurons and glial cells, forming a dynamic scaffold that controls cell differentiation and neuroplasticity (Dzyubenko and Hermann, 2023; Faissner and Reinhard, 2015). To examine cellular composition and phenotypic changes induced by heartbeat sound, we performed single-cell RNA sequencing (scRNAseq). In astrocyte-enriched organoids, heartbeat sound stimulation increased glutamatergic neuron differentiation (23.7 % vs 13.5 % of total cell counts, heartbeat vs control) (Fig. 2A), while no significant cell composition changes were observed in non-enriched organoids (**Fig. S5A**).

**Figure 2.**
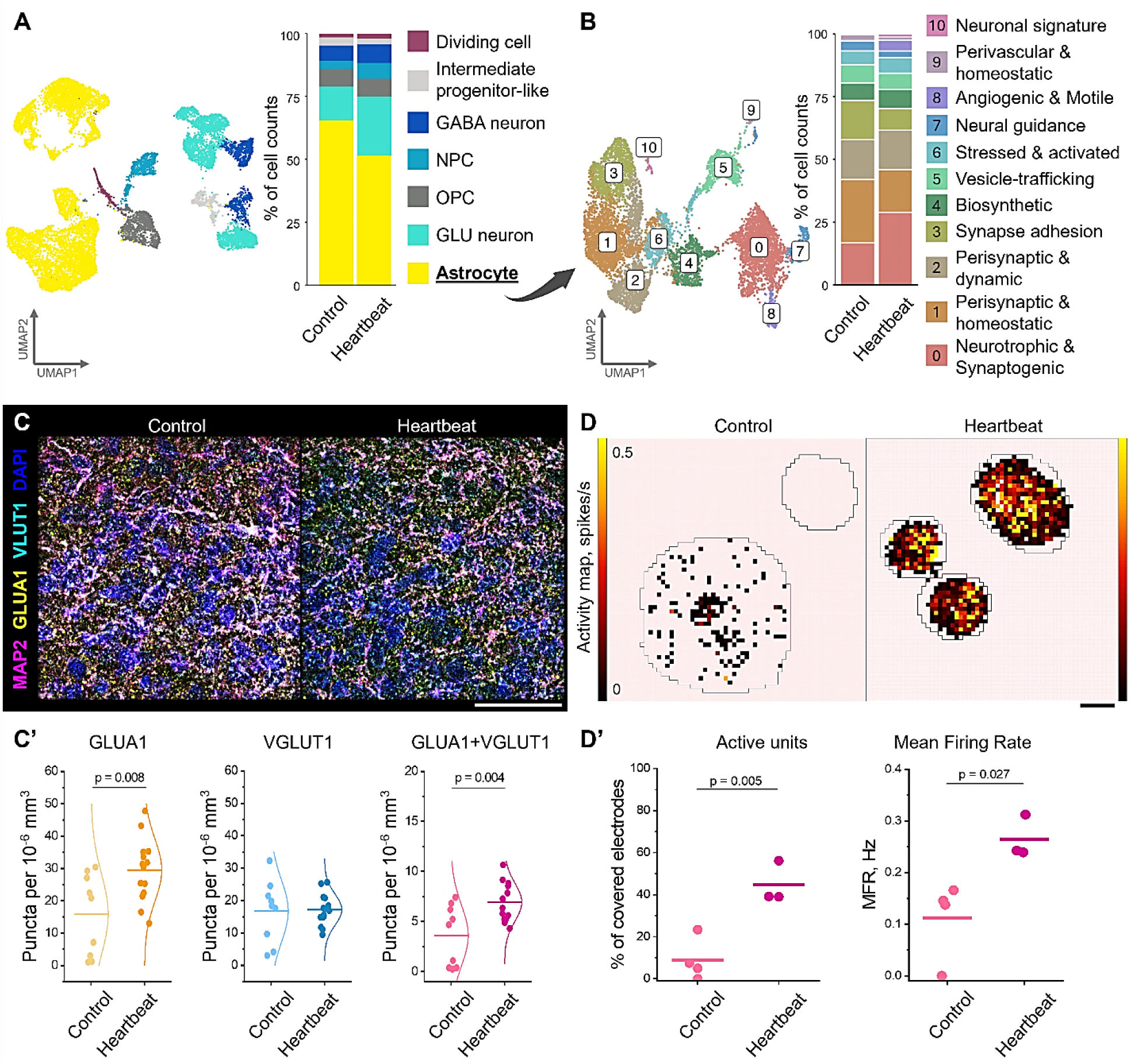
Stimulation with heartbeat sound promotes neural differentiation, synaptogenesis, and spontaneous activity. **(A)** Transcriptomic UMAP map generated from scRNAseq data shows key cellular lineages in astrocyte-enriched organoids. Cell type percentages are shown as stacked bar charts. (**B**) Astrocyte subtypes revealed by cluster analysis of astrocytic transcriptomes (see **Fig. S6** and **Supplementary Table 2** for functional interpretation). Astrocyte subtype percentages are shown as stacked bar charts. (**C**) Immunohistochemical analysis of synaptic markers VGLUT1 (cyan) and AMPA receptor subunit GLUA1 (yellow). Neuronal dendrites are labeled with MAP2 (magenta), and nuclei (blue) are counterstained with DAPI. Scale bar, 50 µm. (**C’**) Quantifications of GLUA1, VGLUT1 synaptic puncta, and their overlaps. (n=9). (**D**) Spontaneous neuronal activity in astrocyte-enriched organoids is shown by high-density multi-electrode array (HD-MEA) activity maps. Scale bar, 500 µm. (**D’**) Quantifications of active units and their mean firing rates (n=3-4). (**C’, D’**) Statistical significance was assessed by unpaired two-tailed t-tests.

A deeper analysis of astrocytic subpopulations by unsupervised clustering (**Fig. S6** and **Supplementary Table 2**) revealed 11 distinct phenotypes (Fig. 2B) in astrocyte-enriched and 6 phenotypes in non-enriched organoids. Among the 11 phenotypes in astrocyte-enriched organoids, heartbeat sound stimulation promoted the cluster of neurotrophic and synaptogenic astrocytes (28.5 % vs 16.1 % of astrocyte cell counts, heartbeat vs control). Organoids without astrocyte enrichment showed no major changes in astrocytic subpopulations (**Fig. S5B**).

### Heartbeat promotes synaptogenesis and activity in astrocyte-enriched organoids

Synaptogenic astrocyte differentiation revealed by scRNAseq was further supported by the upregulation of synaptic proteins detected by LC-MS/MS proteomics. Among the 137 synapse-associated proteins that were upregulated by heartbeat sound stimulation in astrocyte-enriched organoids (**Fig. S7**), we found multiple synaptic adhesion molecules and astrocyte-derived synapse maturation factors including Glypican 4 (GPC4).

Synaptic marker immunolabelling confirmed the increased synaptogenesis in astrocyte-enriched organoids following heartbeat sound stimulation (**Fig. 2C, C’**). Quantifications of postsynaptic puncta containing glutamate receptor subunit 1 (GLUA1), presynaptic vesicular glutamate transporter 1 (VGLUT1), and their overlaps indicated the formation of complete synapses. In cortical organoids without astrocyte enrichment, heartbeat stimulated GLUA1 clustering but not complete synapse formation (**Fig. S8**).

Moreover, high-density multielectrode array (HD-MEA) recordings showed significantly higher spontaneous neuronal activity in heartbeat-stimulated astrocyte-enriched organoids (**Fig. 2D, D’**). While the non-stimulated organoids exhibited little to no spontaneous electrical activity after 31 days of maturation, in agreement with previous reports (Alam El Din et al., 2025; Trujillo et al., 2019), the heartbeat-stimulated organoids demonstrated significantly higher proportion of active units and increased mean firing rates. These results indicate that stimulation with heartbeat sound promotes functional maturation in astrocyte-enriched organoids.

### Heartbeat sound stimulates astrocytic PIEZO receptors

Because sound waves propagate as mechanical deformations in a medium, we hypothesized that the observed effects of heartbeat sound stimulation on organoid maturation can be mediated by mechanoreceptors. scRNAseq data demonstrated that mechanosensitive PIEZO1 and PIEZO2 ion channels (Coste et al., 2010) are selectively expressed by astrocytes and dividing cells (**Fig. 3A**). Because dividing cells are a minor cell population (comprising < 2 % of total cells), we further investigated mechanotransduction of heartbeat sound in astrocytes.

**Figure 3.**
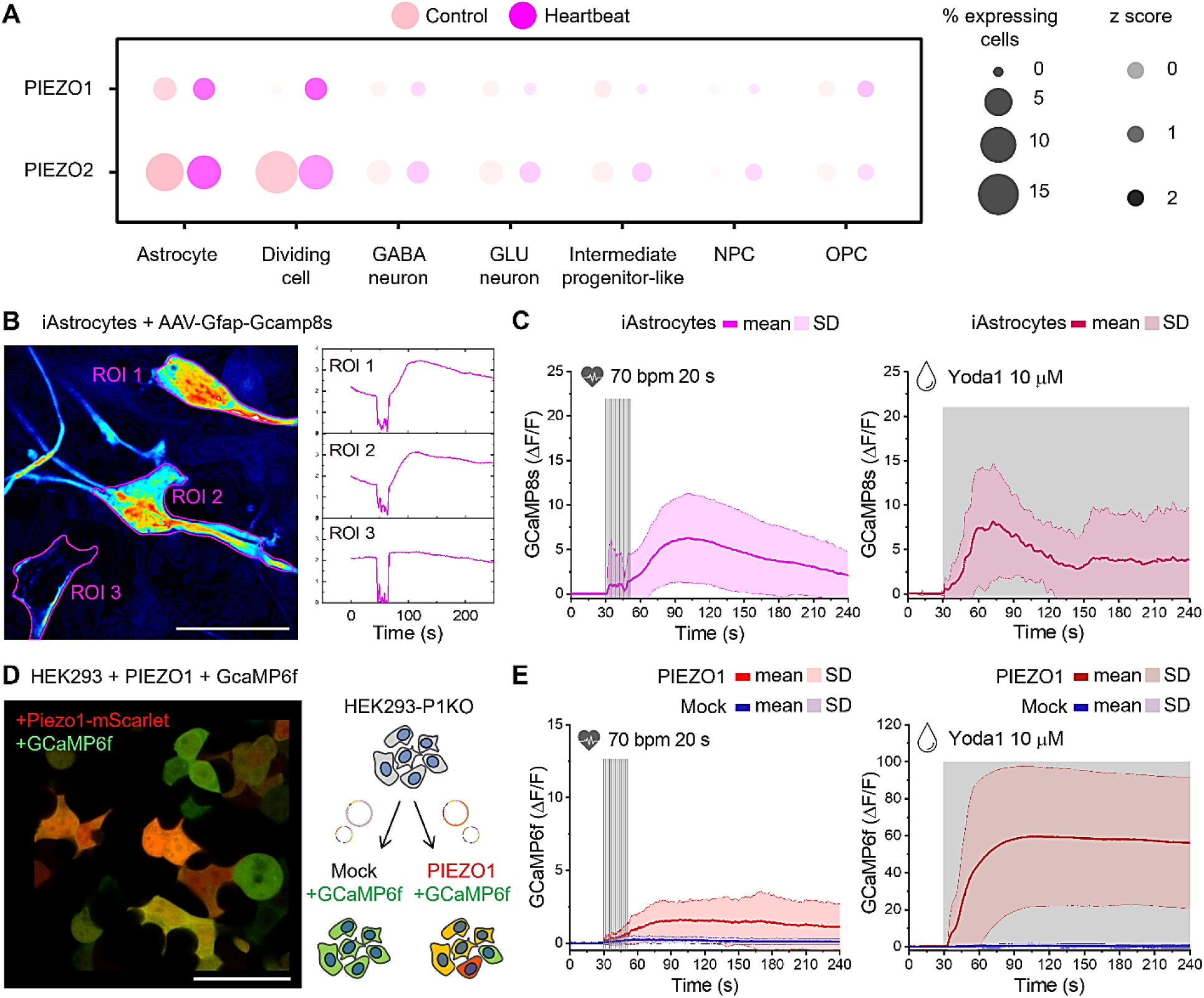
Astrocytic PIEZO ion channels mediate calcium responses evoked by heartbeat. **(A)** Dot plots generated from scRNAseq data show cell-specific expression of mechanosensitive PIEZO channels in astrocyte-enriched organoids. (**B**) Time-lapse t-projection and fluorescence intensity profiles show heartbeat sound-evoked Ca^2+^ responses in human iAstrocytes at 28 days of maturation. Scale bar, 50 µm. (**C**) Averaged Ca^2+^ responses evoked in iAstrocytes (magenta) by heartbeat sound stimulation and 10 µM Yoda1 application (20-25 cells, n=6). (**D** HEK293-P1KO cells were co-transfected with the GCaMP6f Ca^2+^ indicator and PIEZO1-mScarlet or Mock vectors. (**E**) Averaged Ca^2+^ responses evoked by heartbeat sound stimulation and 10 µM Yoda1 application in HEK cells with (red) and without (blue) PIEZO1 (23-39 cells, n=5). Scale bars, 50 µm.

To perform simultaneous acoustic stimulation with heartbeat-like sound and Ca^2+^ imaging, we used a custom-built device (**Fig. S9**), adapted from (Kumeta et al., 2025). Ca^2+^ imaging in maturated iAstrocytes revealed robust responses to both short (20 s) heartbeat sound stimulation (**Table 1**) and bath application of selective PIEZO1 agonist Yoda1 (Evans et al., 2018) (Fig. 3B, C). Notably, the undifferentiated NPCs showed only minimally detectable responses to both heartbeat sound and Yoda1, indicating that the heartbeat sensitivity is a feature of maturating astrocytes (**Fig. S10**).

To test whether heartbeat-like sound stimulation selectively activates PIEZO1 mechanosensitive ion channels, we performed Ca^2+^ imaging in a human embryonic kidney (HEK293) cell model lacking endogenous PIEZO1 expression (HEK293-P1KO) (Dubin et al., 2017). PIEZO1 expression was induced using a plasmid vector with a fluorescent reporter mScarlet to identify transfected cells (Fig. 3D). Both short (20 s) stimulation with heartbeat sound and bath application of Yoda1 elicited sustained Ca^2+^ increase in PIEZO1-expressing cells, whereas mock-transfected controls remained inactive (Fig. 3E). Notably, co-application of heartbeat stimulation with a non-activating Yoda1 concentration (0,01 µM; IC₅₀ in HEK cells ∼1.3 µM) amplified the Ca^2+^ response, while controls remained silent, confirming the specificity of PIEZO1 activation by heartbeat sound (**Fig. S11**).

### PIEZO1 activation triggers ECM expression in mouse astrocytes

To investigate the downstream effects of astrocytic PIEZO activation by low-frequency sound stimulation, we applied short 1 s sound bursts with a dominant frequency of 150 Hz (**<u>Fig.</u> <u>S12A</u>**) in primary mouse astrocytes. This mode of PIEZO1 activation evoked similar responses as human heartbeat sound stimulation in HEK cells (**Fig. S12B**). In mouse astrocytes, short sound bursts evoked more robust responses than human heartbeat sound (**Fig. 4A, B** and **Fig. S12C-F**), making this PIEZO1 activation regime preferable for downstream signaling studies.

**Figure 4.**
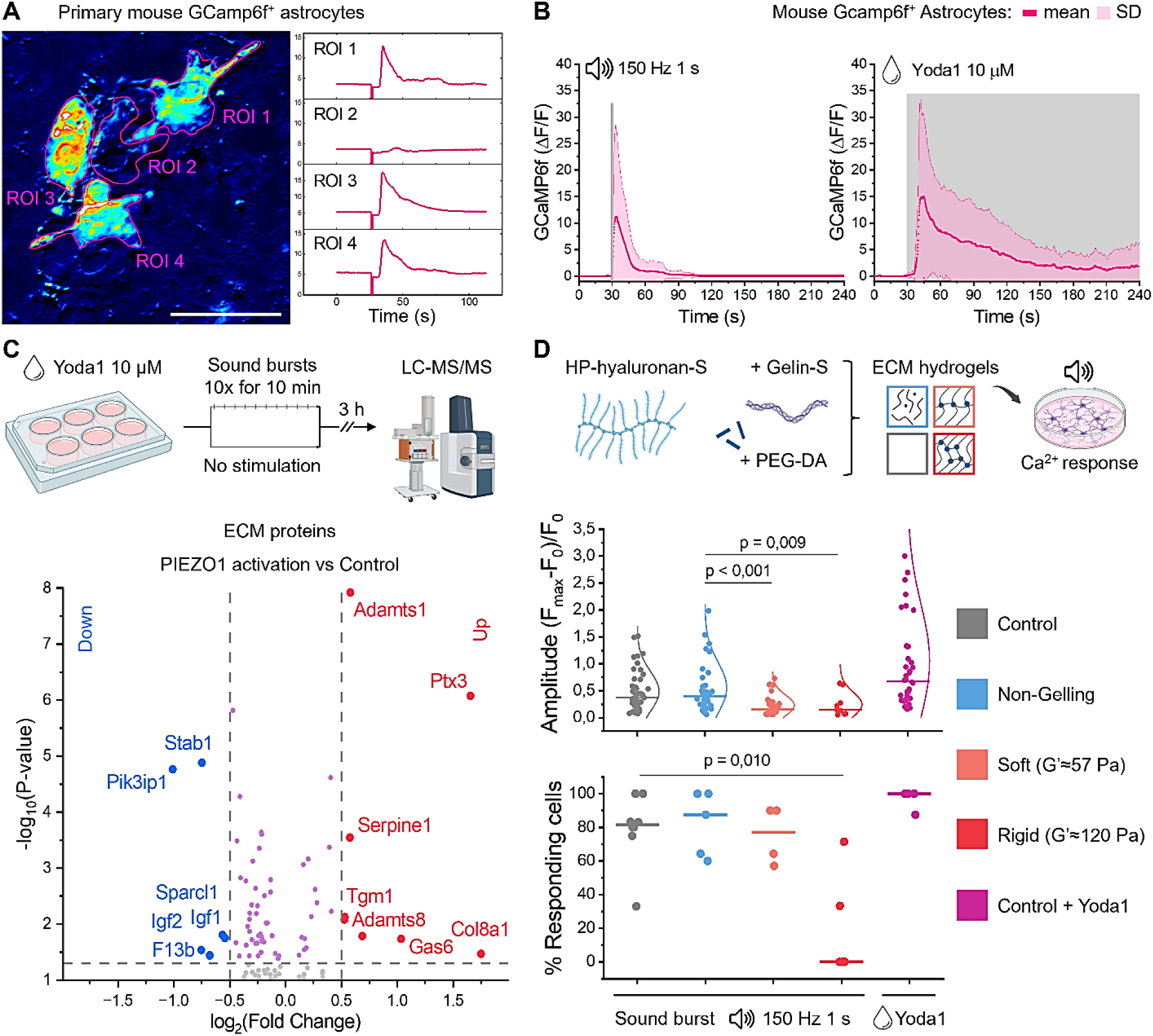
PIEZO1 activation and its downstream signaling in primary mouse astrocytes. **(A)** Time-lapse t-projection and fluorescence intensity profiles show heartbeat-evoked Ca^2+^ responses in mouse primary astrocytes. Scale bar, 50 µm. (**B**) Averaged Ca^2+^ responses evoked in astrocytes by low-frequency sound bursts and 10 µM Yoda1. (38-42 cells, n=8). (**C**) LC-MS/MS measurements were performed 3 hours after PIEZO1 activation in mouse astrocytes. Volcano plot shows key ECM protein regulations induced by PIEZO1 activation, compared versus control. (n=5). For a full list of regulated proteins see **Fig. S13**. (**D**) Ca^2+^ responses to low-frequency sound bursts are quantified in astrocytes covered with biomimetic ECM hydrogels with variable stiffness. (n=4). Intergroup differences were analyzed by one-way ANOVA followed by t-tests.

To reveal protein expression changes induced by astrocytic PIEZO1 activation, we performed LC-MS/MS proteomics following a 3-hour incubation with 10 µM Yoda1 combined with ten acoustic bursts delivered during the first 10 minutes (Fig. 4C). Thereby induced astrocytic PIEZO1 activation increased the expression of ECM scaffolding components including the network-forming collagen COL8A1, pentraxin PTX3, ECM-remodeling enzymes including matrix metalloproteinases with thrombospondin motifs ADAMTS1 and ADAMTS8, and ECM-crosslinking transglutaminase TGM1 (Fig. 4C and **Fig. S13A**). Intracellularly, PIEZO1 activation induced Ca^2+^ response and inflammation-associated transcriptional factor expression (**Fig. S13B**).

### Biomimetic ECM hydrogels attenuate PIEZO1 activation by low-frequency sound

Because the extended activation of PIEZO1 in mouse astrocytes stimulated stress-associated pathways, we hypothesized that the brain ECM can modulate cellular responses to mechanical stimulation such as low-frequency sound. To test this hypothesis, we covered the GCaMP6f-expressing primary mouse astrocytes with ECM-mimetic hydrogels with tunable stiffness corresponding to soft and physiologically stiff brain ECM (**Fig. 4D**). Under 150 Hz sound burst stimulation, the hydrogels significantly attenuated PIEZO activation, indicated by decreased amplitude of Ca^2+^ responses and reduced fraction of responding astrocytes.

## DISCUSSION

Our findings demonstrate a novel mechanism for activating ECM synthesis and neural differentiation via mechanotransduction of physiological sound stimuli. Heartbeat sound stimulation of astrocytic PIEZO ion channels promoted the differentiation of neurotrophic astrocytes and glutamatergic neurons. Neural cell differentiation was accompanied by coordinated expansion of ECS and ECM synthesis, which supported functional neuronal maturation in human cortical organoids. This evidence suggests interoception of the internal acoustic environment as an underappreciated driver of human brain development.

We show that the neurodevelopment and maturation promoting effects of heartbeat sound stimulation in human brain organoids are astrocyte dependent. Heartbeat sound induced ECM synthesis, neural differentiation, synapse formation, and spontaneous neuronal activity exclusively in the astrocyte-enriched organoids. These findings align with the established role of astrocytes in ECM remodeling and regulation of neuroplasticity (Dzyubenko and Hermann, 2023). Consistent with the function of PIEZO ion channels in mouse hippocampal neurogenesis (Chi et al., 2022), we found that activation of astrocytic PIEZO channels by heartbeat sound promoted a neurotrophic and synaptogenic astrocyte phenotype. Given that our scRNAseq data demonstrated predominant PIEZO expression in astrocytic lineages, these results suggest that astrocytes are key interoceptors sensing and transducing internal acoustic stimuli during human neurodevelopment.

This finding was enabled by our astrocyte enrichment protocol, which achieves a physiological 1:1 astrocyte-to-neuron ratio (von Bartheld et al., 2016). Delayed astrocytic differentiation and maturation in conventional human cortical organoids (Sloan et al., 2017) is a limitation for investigating astrocyte-mediated mechanisms. Our approach supports the early emergence of mature, ramified astrocytes. Furthermore, single-cell transcriptomics revealed diverse astrocyte subtypes including perisynaptic and regulatory phenotypes, establishing this model as a powerful tool for investigating astrocyte-neuronal interactions and their contributions to human cortical development.

We demonstrate that PIEZO channels are activated by low-frequency sound stimulation resembling the acoustic environment *in utero* (Gerhardt et al., 1990; Vince et al., 1982). While brief 150 Hz sound bursts elicited transient PIEZO activation, rhythmic heartbeat sound with the dominant frequency of 100 Hz induced sustained intracellular Ca^2+^ elevations. Continuous stimulation with alternating pressure beyond 10 Hz has been shown to rapidly deactivate PIEZO1 (Lewis et al., 2017). However, that effect was observed during prolonged sinusoidal stimulation for 4 seconds, whereas the individual heartbeat pulses used here did not exceed 0.15 seconds, verified by hydrophone recordings. Consistent with previous studies showing neuronal activation in response to repetitive mechanostimulation *in vitro* (Nikolaev et al., 2015) and *in vivo* (Jammal Salameh et al., 2024), we demonstrate that rhythmic heartbeat sound and consecutive sound bursts robustly activate mechanotransduction in astrocytes. Together with proteomic evidence for upregulated Ca^2+^ responsive transcriptional factors and ECM expression, these data indicate that PIEZO activation by heartbeat induces Ca^2+^-dependent ECM synthesis in astrocytes.

Astrocyte-dependent ECM synthesis and ECS expansion facilitated functional maturation of heartbeat-stimulated human brain organoids. While the ECS provides the diffusion routes for ions and metabolites essential for neuronal activity (Sykova and Nicholson, 2008), uncontrolled ECS enlargement can create “dead-space” microdomains that impair rather than facilitate diffusion and signaling (Hrabetová et al., 2003). Stabilization by ECM is therefore crucial, with the core scaffolding components providing a macromolecular network that binds regulatory molecules (Fawcett et al., 2022), mediates neuron-glia interactions (Dzyubenko and Hermann, 2023), and controls neuronal plasticity (Dityatev et al., 2010). Thus, the coordinated increase in ECM synthesis and ECS volume established a homeostatic environment that supported glutamatergic neuron differentiation, synapse formation, and neuronal activity following heartbeat sound stimulation in astrocyte-enriched cortical organoids.

In addition to the maturation-associated benefits of astrocytic mechanostimulation, a prolonged PIEZO activation also induced inflammatory signaling. In this context, our experiments with biomimetic hydrogels indicate a mechanoprotective role of the brain ECM. By attenuating PIEZO activation, physiologically stiff ECM buffers astrocytic responses to mechanical stimulation. This dual role of facilitating supportive signaling while preventing excessive activation suggests ECM is a homeostatic regulator of mechanotransduction in the developing brain.

Our findings open an intriguing direction for developing ECM-promoting therapies in multiple neurological disorders that are associated with ECM degradation, including stroke (Dzyubenko et al., 2023) and Alzheimer’s disease (Crapser et al., 2020). Strategies to restore ECM, particularly neuron-supportive components, remain elusive. We show that these components can be stimulated by low-frequency sound through astrocytic PIEZO activation. While focused ultrasound has been explored for neuromodulation (Kosnoff et al., 2024), its clinical translation has been limited by uncertain mechanisms. By revealing how astrocytes respond to low-frequency sound, this work suggests a promising approach for harnessing astrocyte-mediated ECM remodeling for neurological recovery.

## ACKNOWLEDGEMENTS

This work was supported by the German Research Foundation (DFG, project 467228103 to ED; projects 504501554, 466687329 (SP4), 539301313, and 449437943 (A3,Z1) to DRE; projects 405358801/428817542 (within FOR2879), 449437943 (within TRR332) and 514990328 to DMH). FR acknowledges funding from the Melanoma Research Alliance and the Wolfgang & Gertrud Boettcher Foundation. OS acknowledges funding from the ERA-NET NEURON initiative (project JTC2024 - Brain-Body: BIO-PREPA). We would like to also acknowledge the Center for Advanced Imaging (CAi) at Heinrich Heine University and especially Prof. Stefanie Weidtkamp-Peters and Dr. Sebastian Haensch for the expert help with 3D STED imaging. For the maintenance of confocal and 2P microscopes at the imaging center Essen (IMCES) facility, the authors thank Dr. Anthony Squire, Dr. Jan-Hagen Krohn, and Dr. Anja Hasenberg. For support with HD-MEA electrophysiology, we thank Dr. Elisa Monz (3Brain AG).

## METHODS

### EXPERIMENTAL MODEL

#### Human cortical organoids and induced astrocytes

Neural progenitor cells (NPCs) were differentiated from a highly characterized female control human induced pluripotent stem cell line (SCTi003-A). hIPSCs were plated onto low-adherence AggreWell 800 plates in STEMdiff SMADi neural induction medium and 10µM of Y-27632. During the next 5 days, half of the medium was replaced daily. On day 6, embryonic bodies were replated on a Matrigel coated 6-well plates and full-medium changes were performed daily. On day 12, neural rosette selection was performed using STEMdiff Neural Rosette Selection Reagent for 1,5 hours. After re-plating, daily full-medium changes were performed until the cells formed a monolayer and were ready for passaging. The resulting NPCs can be maintained further with STEMdiff neural progenitor medium.

For quality control and as a shortcut in time-sensitive experiments, ready-to-use NPCs differentiated from the same hIPSC control line (SCTi003-A) were used. Before further differentiation, NPCs were expanded on Matrigel coated 6 well plates in STEMdiff neural progenitor medium, during a maximum of 3 passages, using Accutase enzymatic solution for cell dissociation. NPC cell bank was regularly checked upon each passage for characteristic morphology and marker expression as SOX2 and Nestin, and the absence of OCT4 by immunofluorescence.

Human induced astrocytes (iAstrocytes) were differentiated from NPCs for 3 weeks using STEMdiff astrocyte differentiation medium and maturated using serum-free STEMdif astrocyte maturation medium. iAstrocytes were grown on Matrigel-coated 6-well plates and passaged weekly using Accutase.

For generating human cortical organoids, NPCs were dissociated with Accutase and plated on low-adherence AggreWell 800 plates at 0.2 × 10^6 cells/well. To generate astrocyte-enriched organoids, iAstrocytes at 1 week of maturation were co-seeded together with the NPCs at a 2:1 ratio. Cortical organoid differentiation and maturation was achieved as described by (Birey et al., 2017), with minor modifications. In brief, cortical differentiation was induced using the STEMdiff dorsal forebrain organoid expansion medium containing 20 ng/ml EGF and 20 ng/ml FGF2. After 14 days of differentiation, medium was replaced with the STEMdiff dorsal forebrain organoid differentiation medium containing 20 ng/ml BDNF and 20 ng/ml NT3 for additional 21 days. During these first 35 days, medium was replaced every 2 days. For further maturation, the organoids were maintained in STEMdiff neural organoid maintenance medium without differentiation factors. For analysis, the organoids were transferred to imaging dishes at D45 (31 days of maturation).

#### HEK293 cells with PIEZO1 knockout (HEK293-P1KO)

HEK293 cells lacking PIEZO1 (HEK293-P1KO) were maintained under standard cultivation conditions in DMEM containing 25 mM glucose, 10% fetal bovine serum, and 1% penicillin-streptomycin. The HEK293-P1KO cell line was kindly provided by Ardem Patapoutian’s laboratory (Dubin et al., 2017).

#### Primary mouse astrocytes

Mouse astrocyte cultures were prepared from early postnatal Ai95D (Rosa-CAG-LSL-GCaMP6f) mice as described previously (Borbor et al., 2023; Gottschling et al., 2016). Experimental procedures were approved by the local government (Landesamt für Natur, Umwelt und Verbraucherschutz, Recklinghausen) and conducted according to the European Union directive 2010/63/EU and local institutional guidelines. Ai95D mice were kept in groups of 3-5 animals per cage with inverse 12 h light-dark cycle and *ad libitum* access to food and water.

Cortical mixed glial cultures were obtained from newborn (P1) male and female Ai95 mice. Astrocytic monolayers were purified by cultivation in a selective medium (DMEM with 25 mM glucose, 10% fetal bovine serum, and 1% penicillin-streptomycin) for 7 days. Non-astrocytic cells were removed by shaking (2 hours on a rotary shaker, 150 rpm) and 20 µM cytosine-1-β-D-arabinofuranoside (AraC) application for 24 hours. Purified astrocytes were sub-cultivated in the defined maintenance medium (Neurobasal supplemented with 2% B27, 1 mM L-glutamine, and 1% penicillin-streptomycin) on glass-bottom imaging dishes (µ-Dish 35 mm #81156, Ibidi, Gräfelfing, Germany) pre-coated with 1 mg/ml poly-D-lysine at 1*10^5 cells per dish density.

For inducing Gcamp6f expression, astrocytic monolayers were incubated with 5 µM of cell-permeant nuclei-targeted TAT-Cre recombinase enzyme (#SCR508, MilliporeSigma, Burlington, USA) in standard cultivation medium for 7 days before sub-cultivation on imaging dishes, as described in (Dzyubenko et al., 2021).

### METHOD DETAILS

#### Acoustic stimulation and hydrophone recordings

For long-term stimulation with recorded human 70 beats per minute heartbeat (**Audio File 1**), culture dishes with organoids were maintained on top of continuously emitting (file played in a loop) loudspeaker (Flatpanel CT-17803, ChiliTec) mounted inside of the cell culture incubator and connected to a HiFi digital amplifier and MP3 player (Flatpanel CT-17803, sikkeby). Heartbeat sound levels were adjusted to physiological sound levels (**Table 1**) using the precision hydrophone (8103, HBK) measurements in cell culture medium.

For short-term acoustic stimulation during calcium imaging, 20 seconds of heartbeat (**Audio File 1**) or 1 s sound burst (**Audio File 2**) was played using a custom-made direct stimulation device (Fig. 4B) constructed similar to (Kumeta et al., 2025). In brief, a miniature loudspeaker (8 Ohm, 1 Watt) was attached to the lid of an imaging dish used for confocal microscopy. To achieve the direct transfer of acoustic waves produced by the loudspeaker, a coupler (the red cap of a 15 ml falcon tube) was glued to the loudspeaker membrane. The recorded sound files (**Audio File 1** or **Audio File 2**) were played via an MP3 player using the trigger after the baseline calcium measurements were performed. Sound pressure levels were adjusted close to maximal sound pressure levels *in utero* using the hydrophone.

#### Calcium imaging

For calcium imaging in HEK293-P1KO cell line, cells were plated on imaging dishes (#81156, Ibidi) and co-transfected with 500 ng of pGP-CMV-GCaMP6f (#40755, Addgene, Watertown, USA) and CMV>mPiezo1:IRES:mScarlet3 (#VB241021-1228apt, Vector Builder Inc, Chicago, USA) or empty plasmid pcDNA3.1 (#138209, Addgene) as control. Transient transfections were performed with lipofectamine 3000 transfection reagent (#L3000015, Thermo Fisher Scientific, Oberhausen, Germany) following the manufacturer’s instructions.

For calcium imaging in human iAstrocytes, the cells were plated on Matrigel coated imaging dishes and the pAAV[Exp]-GFAP(short)>jGCaMP8s:IRES:mScarlet3:WPRE virus was added at a multiplicity of infection (MOI) of 5*10^4 for 3 hours. Calcium imaging was performed 1 week after transduction.

For calcium imaging in NPCs, the cells were plated on Matrigel coated imaging dishes and cultivated until monolayer. Thirty minutes before the scans, the culture medium was replaced with Neurobasal medium supplemented with 2% SM1 neuronal supplement and Fluo-4 AM at a final concentration of 5 µM for 30 minutes. Calcium responses were recorded for 5-6 minutes per region of interest including 1 minute baseline at 37°C using the Leica TCS SP8 confocal microscope with a 20x objective. Single plane 129.42×129.42 µm images (pixel size 253×253 nm) were acquired at 0.188 seconds per frame rate). Calcium spikes were detected as (Fmax-Fmin)/Fmin, using standard imageJ selection and measurement tools and custom-written Matlab scripts (**Supplementary Code 1**).

#### Superresolution 3D STED (SuSHi) and 2P shadow imaging

For visualizing the extracellular environment in living organoids, we adapted the SuSHi technique from (Tonnesen et al., 2018). For stability, the organoids were attached to glass-bottom imaging dishes coated with 1:30 diluted Matrigel for 5-10 hours prior to imaging experiment.

3D STED microscopy was performed from the bottom of the organoid, starting 5 µm above the attachment site to avoid potential artifacts from Matrigel coating. For 3D STED shadow imaging, organoid culture medium was supplemented with 200 µM non-cell permeable calcein. 3D STED microscopy was performed using a Leica 3D gSTED SP8 microscope (93x glycerol immersion objective with motor correction, HC PL APO 93x/1,30 GLYC motCORR STED WHITE, and 75% 3D STED). Motor correction was adjusted for maximum resolution at 10 µm imaging depth. Z-stacks (125×125×15 µm, voxel size 61×61×150 nm) were obtained between 5 and 20 µm from the coverslip. Apparent resolution was estimated by measuring the full width at half-maximum (FWHM) of the smallest visible structures at 10 µm depth (**Supplementary Table 1**).

2P microscopy was performed from the top of the organoid, and resolution at 100 µm depth was estimated by FWHM of the smallest detectable structures (**Supplementary Table 1**). For 2P shadow imaging, organoid culture medium was supplemented with 200 µM non-cell permeable Lucifer Yellow. 2P microscopy was performed using a Leica TCS SP8 (25x HCX IRAPO L 25x water immersion objective, NA 0.95) microscope. ROIs (295.3×295.3×115 µm, voxel size 85×85×699 nm) were scanned as z-stacks. The excitation laser (Titanium:Sapphire Chameleon Vision II) was tuned to 890 nm, and the 15% of 1.7 W output power was used. Emitted fluorescence (500 - 570 nm detection wavelength) was collected using a hybrid detector.

#### Immunolabeling procedures

For immunocytochemistry, samples were fixed with 4% PFA for 10 min (2D cell cultures) or 30 min (organoids) at room temperature (RT). Permeabilization and pre-blocking was applied for 2 hours at RT (2D cells) and 5 hours (organoids) with PBS pH 7.0 containing 0.15% Triton X-100 (2D cells) or 0.3% Triton X-100 (organoids), 10% Chemiblocker and 2% donkey serum. Primary antibodies were applied overnight at 4°C, and secondary antibodies conjugated to fluorescent dyes were used for detection. Nuclei were counterstained with DAPI.

To enable 3D confocal imaging of large organoid volumes, optical index of tissues was matched to the target objective immersion by sequential incubation in 50% and 97% 2,2′-thiodiethanol (TDE) for 20 minutes, followed by mounting in 97% TDE on imaging dishes.

#### Microscopy and immunofluorescence quantifications

Image stacks for marker protein quantification in organoids were acquired using the Leica 3D gSTED SP8 microscope with a 20x Plan-Apochromat objective, numerical aperture (NA) 0.8. ROIs (472.3×472.3×5.5 μm, 124×124×500 nm) were scanned as Z-stacks and converted to maximum intensity projections. Image analysis was performed using standard tools in ImageJ.

High-resolution images for synapse quantification in organoids were acquired using the confocal microscope LSM710 (Carl Zeiss) with a 63x Plan-Apochromat objective (NA 1.4). Three random ROIs (90×90×5.5 μm, voxel size 88×88×457 nm) per organoid were scanned Z-stacks. 3D synapse quantification was performed using the Synapse counter plugin for ImageJ (Dzyubenko et al., 2016)

#### High-density multielectrode (HD-MEA) electrophysiology

Spontaneous activity was recorded using the BioCAM DupleX paired with CorePlate 1W 38/60 MEA plates (3Brain AG). Organoids were plated onto the electrode field, and immobilized for recording with the 3D printed meshes (3Brain GmbH, Lanquart, Switzerland). Before recordings, organoids were equilibrated for 1 hour in the incubator. Network activity was recorded for 5 minutes at 37°C using Brainwave 5 software. The recordings were acquired at 20 kHz sampling rate.

#### Biomimetic ECM hydrogels

Biomimetic hydrogels with adjustable stiffness were produced using the HyStem-HP Hydrogel Kit (Advanced Biomatrix) according to manufacturer’s instructions. Gel stiffness was tuned to 0, 57, and 120 Pa shear storage moduli by adjusting the concentration of crosslinking reagent (PEGDA). Before polymerization, gel components were pre-mixed with the cultivation medium and applied in 250 µl droplets onto astrocytic monolayers. After 2 hours, the gels were completely polymerized, and Ca^2+^ imaging was performed 24 hours later.

#### LC-MS-MS proteomics

Human cortical organoids or mouse primary astrocytes were washed in cold PBS and lysed in 30 µl of lysis buffer (Tris-HCl, 150 mM NaCl, 1% SDS, pH 7.8) supplemented with complete mini EDTA free protease inhibitor. Samples were treated with 2,5 U Benzonase and sonicated in a water bath twice for 5 minutes each, and centrifuged at 5000 g for 10 min to remove cell debris. Protein concentration was determined using a BCA assay.

Tryptic digestion was performed for 16 hours at 37° C using the single-pot, solid-phase-enhanced sample preparation (SP3) protocol (Hughes et al., 2019) with trypsin-to-protein ratio of 1:30. Eluted peptides were dried in an Eppendorf Concentrator Plus (Eppendorf SE, Hamburg, Germany) and stored at minus 80° C until further use. Directly prior to measurement, peptides were reconstituted in 20 µl of 0.1% FA and centrifuged at 16,000 g for 5 minutes to remove remaining beads. The peptide concentration was estimated using a NanoDrop Microvolume Spectrometer (Thermo Fischer Scientific, Waltham, USA) at 205nm.

For human cortical organoids, 50 ng of peptides were loaded onto an Evotip (Evosep Biosystems, Odense, Denmark) following the manufacturer instructions. Peptide separation was performed on an Evosep One (Evosep Biosystems, Odense, Denmark) using the predefined Wisper Zoom 40 SPD method on an Aurora Elite CSI size C18 UHPLC column (15 cm length, 75 um inner diameter, 1.7 µm particle size, 120 Å pore size (IonOpticks, Collingwood, Australia)).

For primary mouse astrocytes, 200 ng of peptides were injected into a Nanoelute2 LC system (Bruker Daltronics, Brillerica, Massachusetts, USA). Peptides were loaded onto a PepMap Neo Trap Cartridge (5 mm length, 300 um inner diameter, 5 um particle size, 100 Å pore size; Thermo Fisher Scientific, Massachusetts, USA) and analyzed using a 50-min chromatography gradient on an Aurora Ultimate CSI 18 capillary column (25 cm length, 75 um inner diameter, 1.7 um particle size, 120 Å pore size (IonOpticks, Collingwood, Australia) consisting of mobile phase, A (0.1% FA in H20) and mobile phase B (0.1% FA in ACN-H2O), ranging from 2-34% B at a flow rate of 300nL/min.

Eluting peptides were injected into an TimsTOF HT mass spectrometer (Bruker Daltronics, Brillerica, Massachusetts, USA), via a CaptiveSpray source at 1600V. The mass spectrometer was operated in data-independent acquisition (DIA) PASEF mode. The accumulation and ramp time for the dual TIMS analyzer were set to 100ms at a ramp rate of 9,42 Hz. Scan ranges were set to m/z 100-1700 for the mass range and 1/k0 0.60-1.60Vs-1cm-2 for the mobility range at both MS levels. For DIA-PASEF, a cycle time of 1.8 seconds was estimated. Within each cycle, a precursor mass range of 400 to 1201 Da and a mobility range of 0.6 to 1.6 1/K0 was fragmented. At MS1 level, one ramp was executed per cycle. For MS/MS measurements 32 MS/MS windows with a mass width of 26.0 Da each and a mass overlap of 1.0 were distributed across the 16 MS/MS ramps.

#### Single-cell RNA sequencing

Single-cell suspensions were obtained by digesting 20 organoids per condition in 1 ml of lysis buffer containing 30U of papain, 13mM of L-cysteine, and 125 U of DNAse1. The lysis buffer was pre-activated for 20 minutes at 37°C, and the organoids were dissociated using a rotary shaker at 500 rpm and 37°C for 45 min and by pipetting up and down every 5 minutes.

In each sample, between 12.000 and 20.000 cells were loaded onto a GEM-X single-cell 3’ chip. Sequencing-library construction was performed by following the GEM-X 3’ assay v4 (10x Genomics) protocol. Sequencing was performed using the Novaseq X plus (Illumina) platform aiming for 30k reads per cell (PE150).

### QUANTIFICATION AND STATISTICAL ANALYSIS

#### ECS volume

Segmentation of cellular and extracellular organoid compartments for ECS volume quantifications in 3D STED and 2P shadow imaging was performed using trainable Weka segmentation in ImageJ. Raw images were preprocessed, non-organoid masks detected by custom plugin and subtracted from the segmented extracellular space (**Fig. S2** and **Supplementary Code 2**).

#### Spontaneous activity

For spike detection, the high-pass filter (100 Hz) was applied, and the sliding window differential threshold algorithm implemented in Brainwave 5 software. A threshold of 9*SD of the noise was used, sliding window length 2ms, and the timestamps were assigned to the negative peak of the action potentials. For spike sorting we used the Legendre moments algorithm for feature extraction paired with a K-means & Silhouette algorithm for clustering. Mean firing frequency of each active unit was calculated for the entire recording time. Only units with a mean firing frequency higher than 0.05 Hz were considered. Bursts were defined as series of 5 consecutive spikes occurring within a 100ms window.

#### LC-MS-MS data quantifications

LC-MS raw data were processed using the direct DIA algorithm in Spectronaut (Version 19.1.240806.626, Biognosys, Schlieren, Switzerland), applying BSG factory settings (Enzyme/ Cleavage rules: Trypsin/P; Fixed modification: Carbamidomethyl (C); Variable modifications: Acetyl (Protein N-term) and Oxidation(M)), against a reviewed human UniProt/Swissprot FASTA database (from 13th of March 2024, containing 20418 target sequences) for human cortical organoids and against a reviewed murine UniProt/Swissprot FASTA database (from 13th of March 2024, containing 17196 target sequences). Quantification was performed at the MS2 level. Imputation was disabled. Cross run normalization was enabled. Normalized protein abundance was used for further downstream analysis, exported and analyzed in R.

Protein abundances were log2 transformed to approach the Gaussian probability distribution. Differentially abundant proteins between multiple predefined phenotypes were determined by analysis of variation (ANOVA). Significantly regulated proteins (p<0.05) were subjected to further analysis. Differentially abundant proteins between pairs of predefined phenotypes were determined by Student’s t-testing. Proteins showing significant differences (p<0.05, fold change > 1.5) were subjected to further analysis. Heatmaps were generated based on hierarchical clustering results. The Pearson correlation was used as a distance metric at the sample and protein level respectively. Ward.D2 linkage was applied. Prior to heatmap generation, log2 protein abundances were centered around the mean per protein for visual purposes. For principal component analysis, a missing data-tolerant nonlinear PCA (NIPALs-PCA) was used.

#### Single-cell RNA sequencing quantifications

Raw sequencing data was processed using CellRanger (version 9.0.1) and aligned to the human reference genome (GRCh38). Initial quality control was performed to retain cells with more than 1000 and less than 8000 detected genes, and less than 10% mitochondrial reads. Doublets were identified and removed using DoubletFinder (rate=8%). After quality filtering, a total of 44,743 cells remained for downstream analysis.

Data integration and clustering were performed using the Seurat package (Hao et al., 2024) in R. The filtered data was processed using NormalizeData, FindVariableFeatures, and ScaleData functions. Dimensionality reduction was first performed using RunPCA. Batch effects between different experimental conditions were then corrected using Harmony (Korsunsky et al., 2019) integration based on the PCA results. Visualization was achieved using RunUMAP, and unsupervised clustering was performed using FindNeighbors and FindClusters based on the Harmony embeddings with dims = 1:20 and resolution = 0.3. Cortical organoid single-cell spaces were manually cured by removing fibroblasts. All violin plots, heatmaps, and feature plots were drawn using standard plotting functions from R, Seurat, and SeuratExtend package (Hua et al., 2025). Astrocyte signature intensities were measured using the Bioconductor package AUCell (Aibar et al., 2017).

#### Statistical analysis of imaging and electrophysiology data

Data were analyzed in OriginPro2025 (OriginLab, Northampton, MA, U.S.A.). The normality of data distributions was verified by Shapiro-Wilk mothod. Differences between group means were analyzed using one or two -way mixed model ANOVA with Tukey’s multiple comparisons adjustment, followed by pairwise posthoc comparisons using t-tests. In all experiments, the significance level was set as α=0.05, and the p<0.05 values indicated significant differences between groups.

**Table 2.**
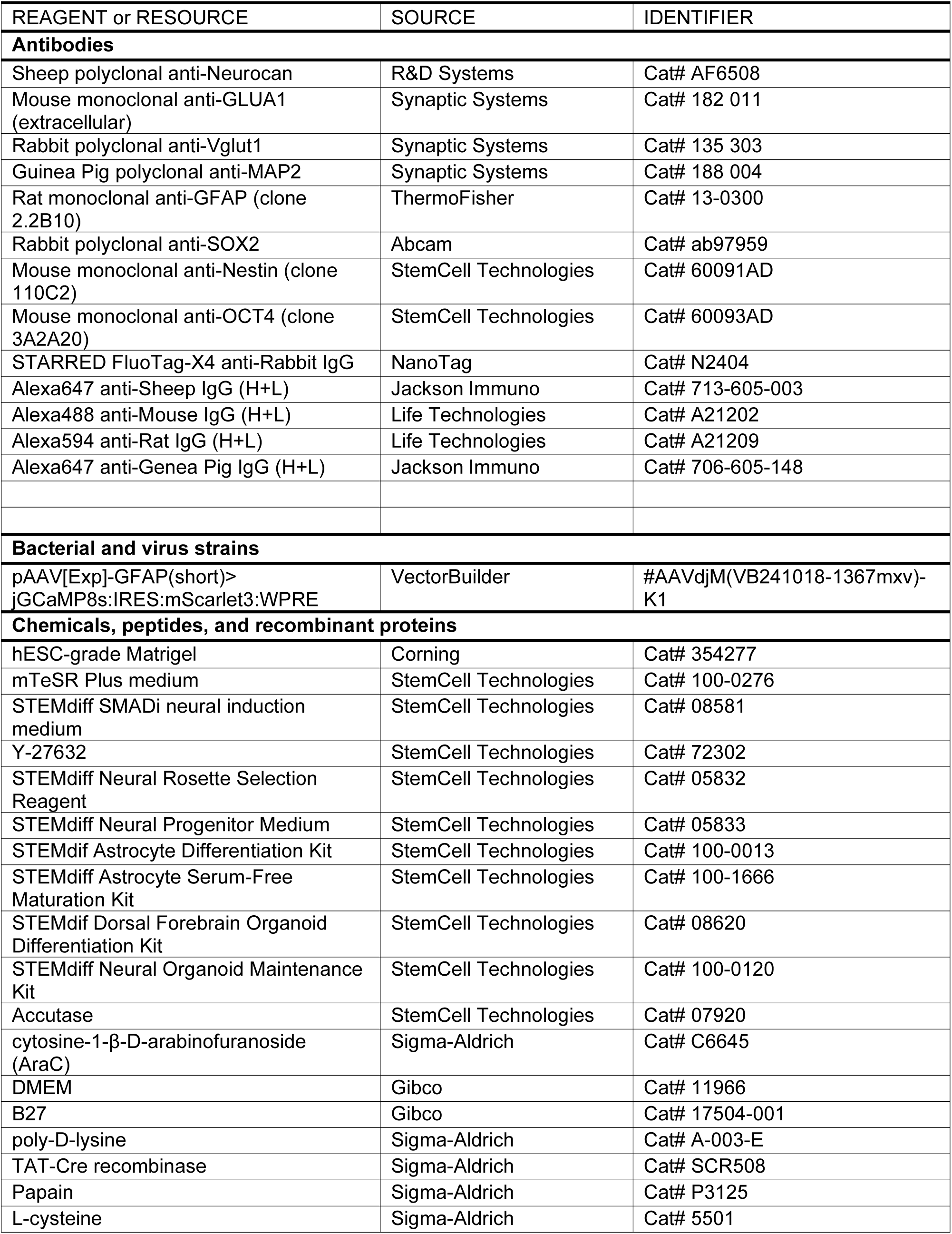
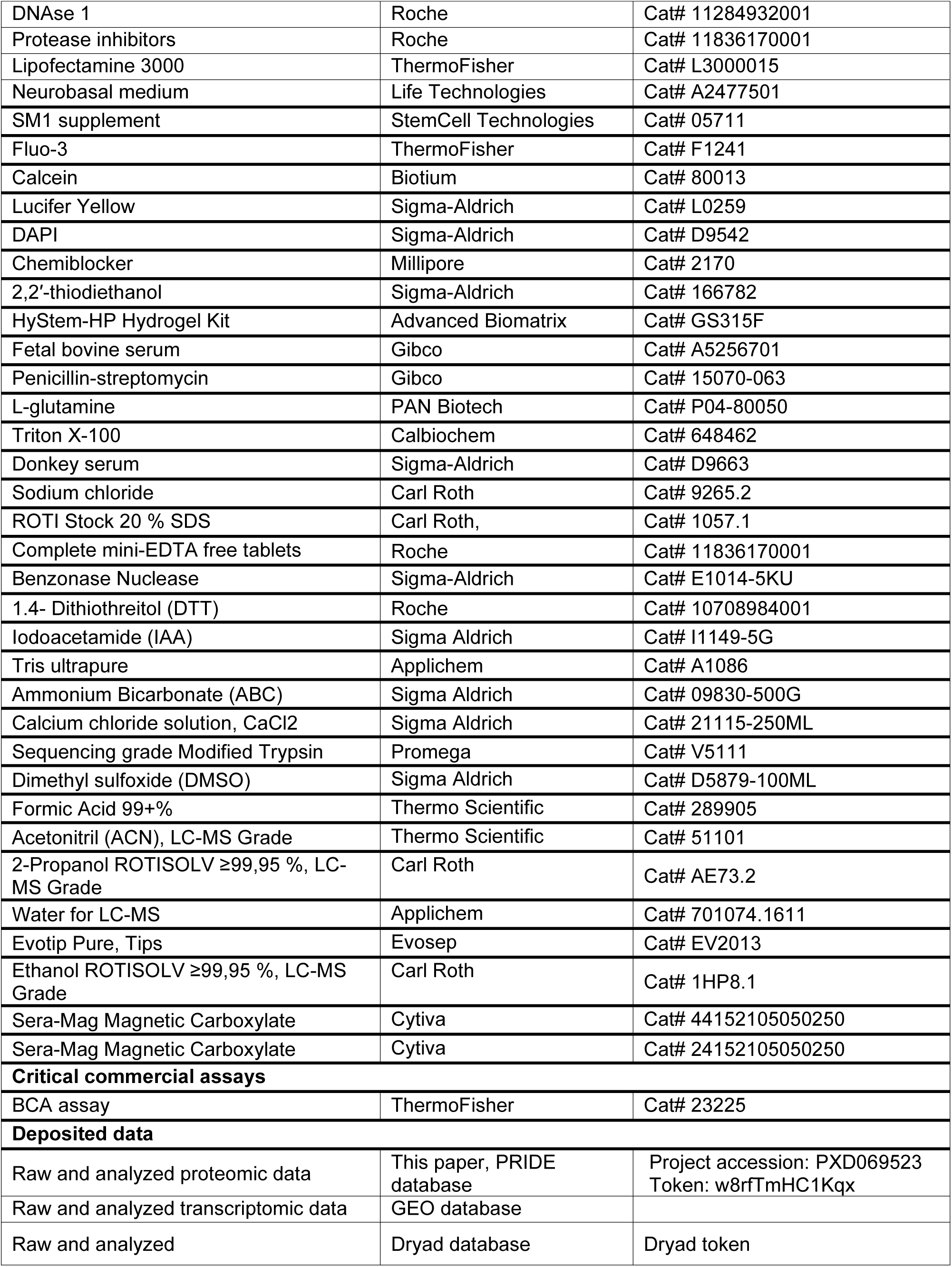
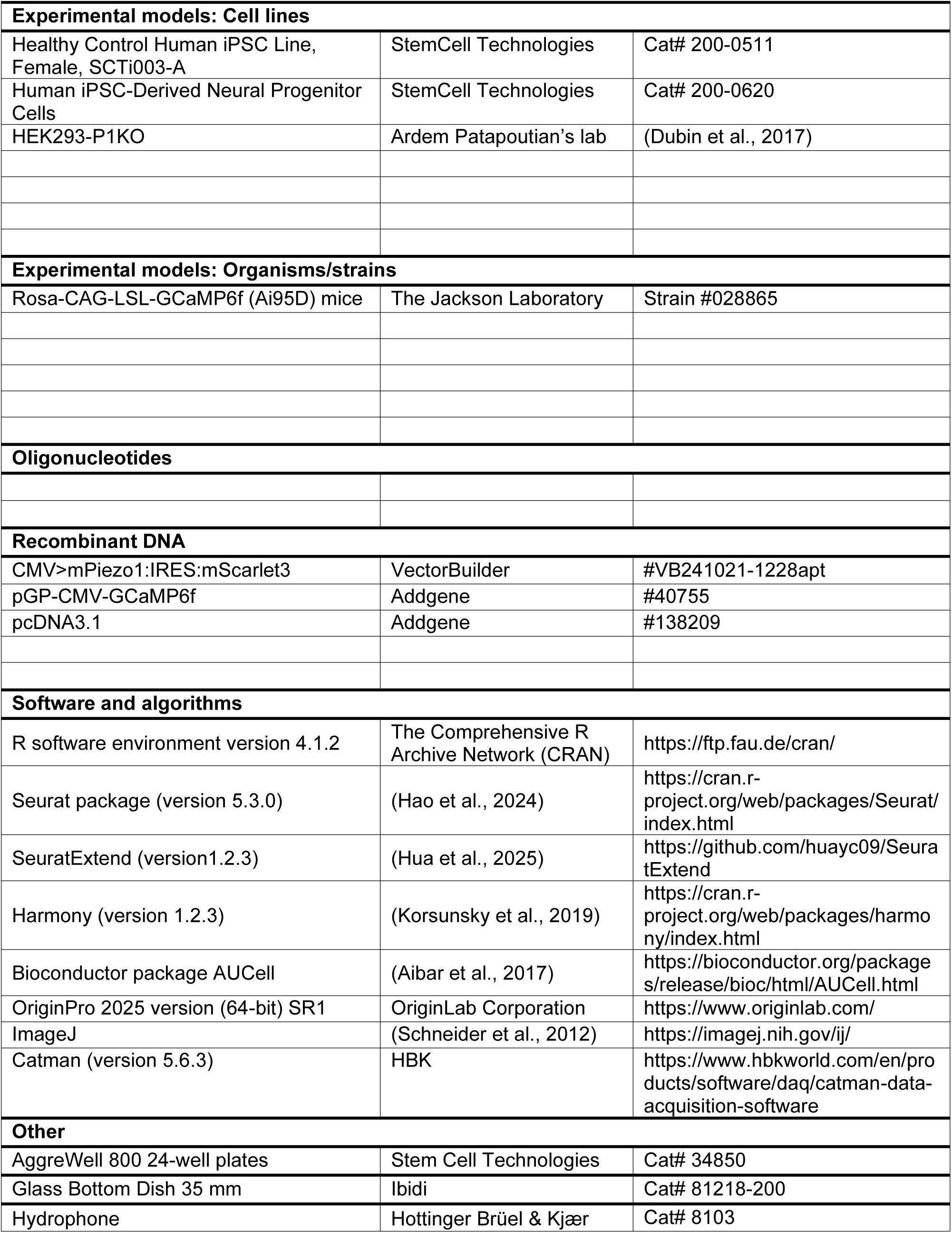
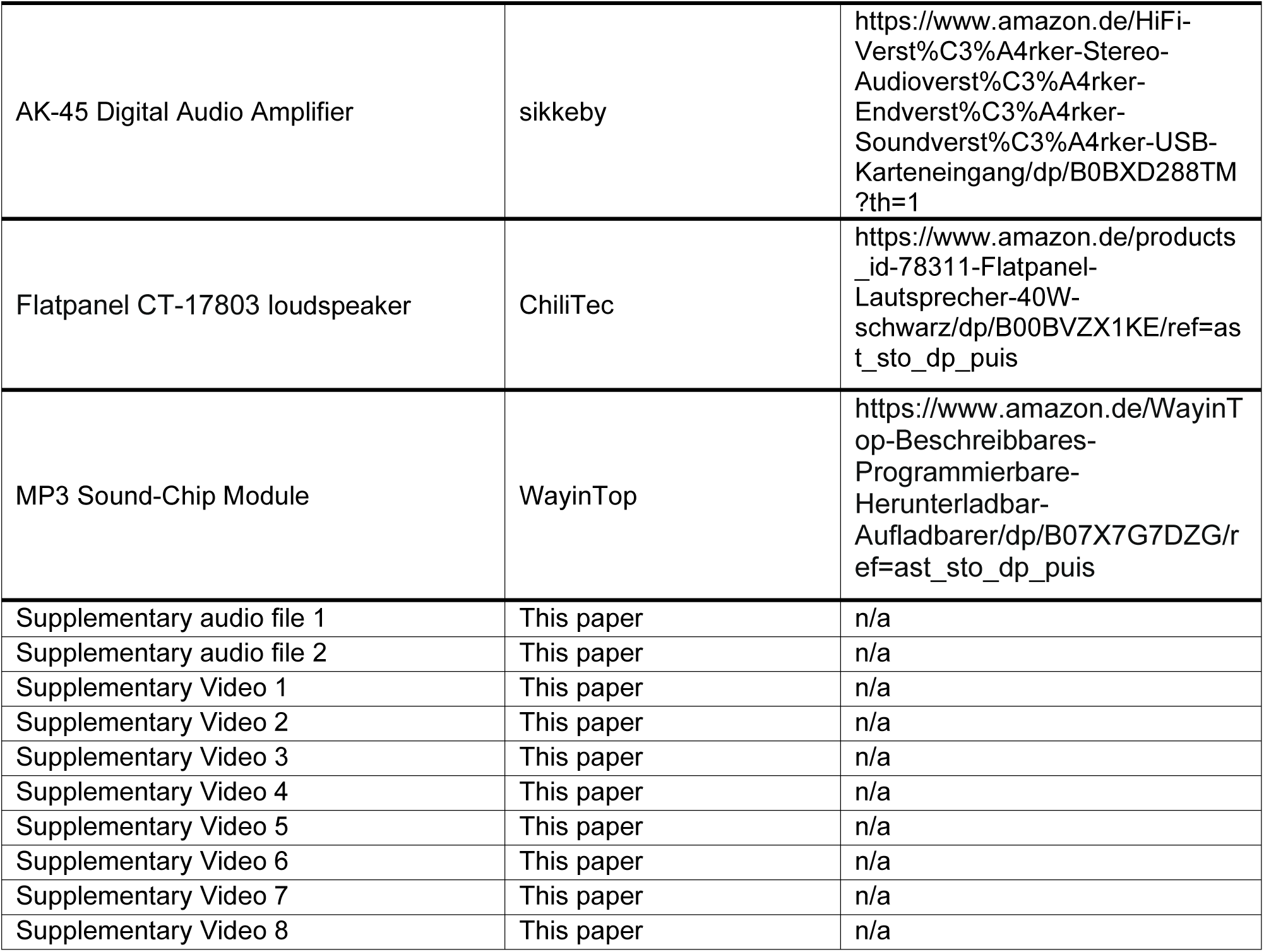
Key resources.

| REAGENT or RESOURCE | SOURCE | IDENTIFIER |
| --- | --- | --- |
| **Antibodies** |  |  |
| Sheep polyclonal anti-Neurocan | R&D Systems | Cat# AF6508 |
| Mouse monoclonal anti-GLUA1 (extracellular) | Synaptic Systems | Cat# 182 011 |
| Rabbit polyclonal anti-Vglut1 | Synaptic Systems | Cat# 135 303 |
| Guinea Pig polyclonal anti-MAP2 | Synaptic Systems | Cat# 188 004 |
| Rat monoclonal anti-GFAP (clone 2.2B10) | ThermoFisher | Cat# 13-0300 |
| Rabbit polyclonal anti-SOX2 | Abcam | Cat# ab97959 |
| Mouse monoclonal anti-Nestin (clone 110C2) | StemCell Technologies | Cat# 60091AD |
| Mouse monoclonal anti-OCT4 (clone 3A2A20) | StemCell Technologies | Cat# 60093AD |
| STARRED FluoTag-X4 anti-Rabbit IgG | NanoTag | Cat# N2404 |
| Alexa647 anti-Sheep IgG (H+L) | Jackson Immuno | Cat# 713-605-003 |
| Alexa488 anti-Mouse IgG (H+L) | Life Technologies | Cat# A21202 |
| Alexa594 anti-Rat IgG (H+L) | Life Technologies | Cat# A21209 |
| Alexa647 anti-Gene Pig IgG (H+L) | Jackson Immuno | Cat# 706-605-148 |
| **Bacterial and virus strains** |  |  |
| pAAV[Exp]-GFAP(short)>jGCaMP8s:IRES:mScarlet3:WPRE | VectorBuilder | #AAVdjM(VB241018-1367mxv)-K1 |
| **Chemicals, peptides, and recombinant proteins** |  |  |
| hESC-grade Matrigel | Corning | Cat# 354277 |
| mTeSR Plus medium | StemCell Technologies | Cat# 100-0276 |
| STEMdiff SMADi neural induction medium | StemCell Technologies | Cat# 08581 |
| Y-27632 | StemCell Technologies | Cat# 72302 |
| STEMdiff Neural Rosette Selection Reagent | StemCell Technologies | Cat# 05832 |
| STEMdiff Neural Progenitor Medium | StemCell Technologies | Cat# 05833 |
| STEMdif Astrocyte Differentiation Kit | StemCell Technologies | Cat# 100-0013 |
| STEMdiff Astrocyte Serum-Free Maturation Kit | StemCell Technologies | Cat# 100-1666 |
| STEMdif Dorsal Forebrain Organoid Differentiation Kit | StemCell Technologies | Cat# 08620 |
| STEMdiff Neural Organoid Maintenance Kit | StemCell Technologies | Cat# 100-0120 |
| Accutase | StemCell Technologies | Cat# 07920 |
| cytosine-1-β-D-arabinofuranoside (AraC) | Sigma-Aldrich | Cat# C6645 |
| DMEM | Gibco | Cat# 11966 |
| B27 | Gibco | Cat# 17504-001 |
| poly-D-lysine | Sigma-Aldrich | Cat# A-003-E |
| TAT-Cre recombinase | Sigma-Aldrich | Cat# SCR508 |
| Papain | Sigma-Aldrich | Cat# P3125 |
| L-cysteine | Sigma-Aldrich | Cat# 5501 |
| DNase 1 | Roche | Cat# 11284932001 |
| Protease inhibitors | Roche | Cat# 11836170001 |
| Lipofectamine 3000 | ThermoFisher | Cat# L3000015 |
| Neurobasal medium | Life Technologies | Cat# A2477501 |
| SM1 supplement | StemCell Technologies | Cat# 05711 |
| Fluo-3 | ThermoFisher | Cat# F1241 |
| Calcein | Biotium | Cat# 80013 |
| Lucifer Yellow | Sigma-Aldrich | Cat# L0259 |
| DAPI | Sigma-Aldrich | Cat# D9542 |
| Chemiblocker | Millipore | Cat# 2170 |
| 2,2'-thiodiethanol | Sigma-Aldrich | Cat# 166782 |
| HyStem-HP Hydrogel Kit | Advanced Biomatrix | Cat# GS315F |
| Fetal bovine serum | Gibco | Cat# A5256701 |
| Penicillin-streptomycin | Gibco | Cat# 15070-063 |
| L-glutamine | PAN Biotech | Cat# P04-80050 |
| Triton X-100 | Calbiochem | Cat# 648462 |
| Donkey serum | Sigma-Aldrich | Cat# D9663 |
| Sodium chloride | Carl Roth | Cat# 9265.2 |
| ROTI Stock 20 % SDS | Carl Roth, | Cat# 1057.1 |
| Complete mini-EDTA free tablets | Roche | Cat# 11836170001 |
| Benzonase Nuclease | Sigma-Aldrich | Cat# E1014-5KU |
| 1,4- Dithiothreitol (DTT) | Roche | Cat# 10708984001 |
| Iodoacetamide (IAA) | Sigma Aldrich | Cat# I1149-5G |
| Tris ultrapure | Applichem | Cat# A1086 |
| Ammonium Bicarbonate (ABC) | Sigma Aldrich | Cat# 09830-500G |
| Calcium chloride solution, CaCl<sub>2</sub> | Sigma Aldrich | Cat# 21115-250ML |
| Sequencing grade Modified Trypsin | Promega | Cat# V5111 |
| Dimethyl sulfoxide (DMSO) | Sigma Aldrich | Cat# D5879-100ML |
| Formic Acid 99+% | Thermo Scientific | Cat# 289905 |
| Acetonitril (ACN), LC-MS Grade | Thermo Scientific | Cat# 51101 |
| 2-Propanol ROTISOLV ≥99,95 %, LC-MS Grade | Carl Roth | Cat# AE73.2 |
| Water for LC-MS | Applichem | Cat# 701074.1611 |
| Evotip Pure, Tips | Evosep | Cat# EV2013 |
| Ethanol ROTISOLV ≥99,95 %, LC-MS Grade | Carl Roth | Cat# 1HP8.1 |
| Sera-Mag Magnetic Carboxylate | Cytiva | Cat# 44152105050250 |
| Sera-Mag Magnetic Carboxylate | Cytiva | Cat# 24152105050250 |
| **Critical commercial assays** |  |  |
| BCA assay | ThermoFisher | Cat# 23225 |
| **Deposited data** |  |  |
| Raw and analyzed proteomic data | This paper, PRIDE database | Project accession: PXD069523<br>Token: w8rfTmHC1Kqx |
| Raw and analyzed transcriptomic data | GEO database |  |
| Raw and analyzed | Dryad database | Dryad token |

| Experimental models: Cell lines |  |  |
| --- | --- | --- |
| Healthy Control Human iPSC Line, Female, SCTi003-A | StemCell Technologies | Cat# 200-0511 |
| Human iPSC-Derived Neural Progenitor Cells | StemCell Technologies | Cat# 200-0620 |
| HEK293-P1KO | Ardem Patapoutian's lab | (Dubin et al., 2017) |
| Experimental models: Organisms/strains |  |  |
| Rosa-CAG-LSL-GCaMP6f (Ai95D) mice | The Jackson Laboratory | Strain #028865 |
| Oligonucleotides |  |  |
| Recombinant DNA |  |  |
| CMV>mPiezo1:IRES:mScarlet3 | VectorBuilder | #VB241021-1228apt |
| pGP-CMV-GCaMP6f | Addgene | #40755 |
| pcDNA3.1 | Addgene | #138209 |
| Software and algorithms |  |  |
| R software environment version 4.1.2 | The Comprehensive R Archive Network (CRAN) | https://ftp.fau.de/cran/ |
| Seurat package (version 5.3.0) | (Hao et al., 2024) | https://cran.r-project.org/web/packages/Seurat/index.html |
| SeuratExtend (version 1.2.3) | (Hua et al., 2025) | https://github.com/huayc09/SeuratExtend |
| Harmony (version 1.2.3) | (Korsunsky et al., 2019) | https://cran.r-project.org/web/packages/harmony/index.html |
| Bioconductor package AUCell | (Aibar et al., 2017) | https://bioconductor.org/packages/release/bioc/html/AUCell.html |
| OriginPro 2025 version (64-bit) SR1 | OriginLab Corporation | https://www.originlab.com/ |
| ImageJ | (Schneider et al., 2012) | https://imagej.nih.gov/ij/ |
| Catman (version 5.6.3) | HBK | https://www.hbkworld.com/en/products/software/daq/catman-data-acquisition-software |
| Other |  |  |
| AggreWell 800 24-well plates | Stem Cell Technologies | Cat# 34850 |
| Glass Bottom Dish 35 mm | Ibidi | Cat# 81218-200 |
| Hydrophone | Hottinger Brüel & Kjær | Cat# 8103 |
| AK-45 Digital Audio Amplifier | sikkeby | https://www.amazon.de/HiFi-Verst%C3%A4rker-Stereo-Audioverst%C3%A4rker-Endverst%C3%A4rker-Soundverst%C3%A4rker-USB-Karteneingang/dp/B0BXD288TM?th=1 |
| Flatpanel CT-17803 loudspeaker | ChiliTec | https://www.amazon.de/products_id-78311-Flatpanel-Lautsprecher-40W-schwarz/dp/B00BVZX1KE/ref=ast_sto_dp_puis |
| MP3 Sound-Chip Module | WayinTop | https://www.amazon.de/WayinTop-Beschreibbares-Programmierbare-Herunterladbar-Aufladbarer/dp/B07X7G7DZG/ref=ast_sto_dp_puis |
| Supplementary audio file 1 | This paper | n/a |
| Supplementary audio file 2 | This paper | n/a |
| Supplementary Video 1 | This paper | n/a |
| Supplementary Video 2 | This paper | n/a |
| Supplementary Video 3 | This paper | n/a |
| Supplementary Video 4 | This paper | n/a |
| Supplementary Video 5 | This paper | n/a |
| Supplementary Video 6 | This paper | n/a |
| Supplementary Video 7 | This paper | n/a |
| Supplementary Video 8 | This paper | n/a |

## SUPPLEMENTARY FILES AND CODE

Supplementary Videos 1-8

Supplementary Codes 1-3

Supplementary audio files 1-2

### Supplementary figures and tables

**Supplementary Table 1.** Shadow imaging resolution.

|  | 3D STED | 2P |
| --- | --- | --- |
| | 10 $\mu\text{m}$ depth | 100 $\mu\text{m}$ depth |
| n | FWHM, nm | FWHM, nm |
| 1 | 150 | 540 |
| 2 | 130 | 460 |
| 3 | 210 | 320 |
| 4 | 130 | 400 |
| 5 | 140 | 590 |
| 6 | 150 | 440 |
| 7 | 120 | 310 |
| 8 | 130 |  |
| 9 | 150 |  |
| 10 | 120 |  |
| 11 | 200 |  |
| 12 | 180 |  |
| 13 | 210 |  |

**Supplementary Table 2.** Astrocyte clusters and their function.

|  | Cluster name | Upregulated genes | Rationale |
| --- | --- | --- | --- |
| **Astrocyte-enriched organoids** |  |  |  |
| 0 | Neurotrophic & Synaptogenic | MDK, IGFBP2, RSPO3, FBLN1, CDH7, TSPAN7, NELL1, ITM2C/BRI2, CLYBL, SPARCL1, WNT7B | Astrocytes enriched for trophic and axon-guidance factors (MDK, IGFBP2, NELL1), Wnt potentiation (RSPO3), ECM assembly (FBLN1), adhesion (CDH7) and EV-linked synapse modulators (TSPAN7), consistent with astrocyte-driven synapse maturation and astro support. These cells likely enhance neurite growth and synaptogenesis while stabilizing the perisynaptic microenvironment. |
| 1 | Perisynaptic & Homeostatic | GJA1, DBI, SCD, TXNIP, MT2A, CPE, PMP2, S100B, EDNRB, AHCYL1, ID2, GABBR2, PCDH9, BCAN | Defined by canonical astrocyte markers (S100B, GJA1, DBI, MT2A, TXNIP) plus BCAN, PCDH9, and GABBR2, this state is related to synapse regulation and homeostatic astrocyte responses to Ca<sup>2+</sup> oscillations |
| 2 | Perisynaptic & Dynamic | DBI, TXNIP, BCAN, AHCYL1, TFPI, WLS, CREB1L, ARID5B, FSTL5, SNCAIP, EDNRB, ID2 | Characterized by Wnt secretion (WLS), synapse interaction adhesion molecules (FSTL5, SNCAIP), and CREB-mediated transcriptional reprogramming, this state combines homeostatic ECM support (BCAN, DBI, TFPI) with enhanced potential for synaptic remodeling and plasticity |
| 3 | Synapse adhesion | PCDH9, ADGRB3, TNIK, GREB1L, PPP2R2B, LRRC4C, MIR99AHG, GLIS3, CTNND2, SNCAIP, FSTL5, NEAT1 | Enrichment of synapse adhesion regulators together with NEAT1, defines a perisynaptic, adhesion-rich astrocyte state that stabilizes and modulates synaptic contacts; reduced ribosomal/EIF1/RACK1 and vimentin indicates a mature, low-biosynthesis, non-reactive profile |
| 4 | Biosynthetic | PMP2, MT2A, SCD, DBI, S100B, RACK1, FTH1, RPLP1, RPLP0, RPL15, RPS28, RPL13A, RPL13, RPS14, RPS8, RPL41 | Marked by ribosomal gene enrichment and reduced developmental adhesion programs (NFIA, ZBTB20, NPAS3, RORA; LSAMP, ADGRV1), this state reflects a translation-high phenotype |
| 5 | Vesicle-<br>trafficking &<br>Synaptogenic | CD9, CDH7, S100A10,<br>S100A6, TSPAN7,<br>NELL2, CAV1,<br>ETNPPL, ANXA2,<br>YBX3, VIM, SPARCL1 | Enrichment of tetraspanins and vesicle-associated genes defines an astrocytic phenotype specialized for exosome- and caveolae-mediated membrane trafficking and intercellular signalling |
| 6 | Stressed &<br>Activated | RACK1, UCHL1,<br>SQSTM1, CEBPB,<br>CCPG1, SARS, DDIT3,<br>TCEA1, HSPA9, EIF1 | This phenotype combines adaptive responses to stress with reduced protein synthesis and upregulated hypoxia-activated genes |
| 7 | Neural<br>patterning &<br>guidance | ZBTB20, EIF4G3,<br>LSAMP, PHACTR1,<br>NFIA, ADGRV1,<br>NPAS3, LRP1B,<br>RORA, NELL1, GLIS3,<br>KALRN | Enrichment of transcriptional factors associated with differentiation, morphogenesis, and adhesion indicates a patterning and contact-guidance astrocytic phenotype |
| 8 | Angiogenic &<br>motile | PTN, NR2F2-AS1,<br>PDE1A, EEF1A1,<br>H3F3A, WNT7B,<br>NR2F2, C1orf61,<br>FABP7, RND3,<br>MARCKSL1 | This cluster combines neurovascular signature (WNT7B, NR2F2, PTN) with a radial glia like expression (FABP7, RND3, MARCKSL1) and Ca<sup>2+</sup> signaling, indicating these astrocytes can migrate and promote vascular development. |
| 9 | Perivascular &<br>homeostatic | S100A6, KCTD1,<br>LHFPL6, NKX2-1,<br>MGP, ATP1A2, EMP2,<br>CALD1, CYSTM1,<br>APOE | This phenotype shows genes related to astrocytic end feet at blood-brain barrier interfaces (ATP1A2, APOE, EMP2, CALD1), secreted matricellular cues (SPARCL1, CD9, S100A10, ANXA2), and ventral telencephalon identity (NKX2-1) |
| 10 | Neuronal<br>signature | GRIA2, DLGAP1,<br>NRXN1, ANK3,<br>MYT1L, KALRN, NAV3 | Canonical neuronal gene expression associated with neurite and synapse development, possibly indicating phagocytic astrocytes contributing to synaptophagy and therefore incorporating neuronal mRNA |

| Cortical organoids without astrocyte enrichment |  |  |  |
| --- | --- | --- | --- |
| 0 | Neurotrophic & Synaptogenic | ETNPPL, PLP1, RSPO3, WNT7B, PHACTR1, RMST, NR2F2-AS1, SPARCL1, ADGRV1, ADAMTSL1, FBLN1, NRXN3, CSRP2, NELL2, AC092957.1 | Astrocytes strongly expressing neurotrophic and synaptogenesis regulators (RSPO3/WNT7B) together with synapse-promoting (SPACRL1) and ECM/adhesion organizers (NRXN3, ADGRV1, ADAMTSL1, FBLN1), consistent with astrocytes that produce trophic cues and actively promote excitatory synaptogenesis. |
| 1 | Homeostatic & Secretory | PTGDS, MT3, C1orf61, CPE, B2M, ID2, CLU, PI15, PCDH9, PPP2R2B | Homeostatic astrocytes enriched with secreted chaperones and neuroprotective signals (CLU, CPE, PTGDS, MT3), while expressing synapse-associated molecules (PCDH9/PPP2R2B) to stabilize and tune astrocyte-neuronal interactions |
| 2 | Synaptogenic & ECM-remodeling | CST3, SPARCL1, S100A10, ADAMTSL1, CD9, TNC, LGALS1, FBLN1, CSRP2, NELL2 | These astrocytes promote and stabilize excitatory synapses (SPARCL1, CSRP2) exporting extracellular vesicles (CD9), and sculpting perisynaptic matrix (CST3, ADAMTSL1, TNC, FBLN1/ADAMTSL1) |
| 3 | Stress-adaptive | FTL, FTH1, CDKN1A, EIF1, RACK1, TKT, RPS27L, RPL19, RPS13, GAS5 | Astrocytes with upregulated DNA repair (CDKN1A, RPS27L), antioxidant response (GAS5, TKT, RACK1, EIF1) and ferroptosis-regulating genes (FTH1, FTL). |
| 4 | Hypoxia-adaptive | IGFBP2, IGFBP5, BNIP3, VEGFA, PGK1, TRIM2, SLC2A1, FAM162A, HILPDA, P4HA1 | These astrocytes show the hypoxia-associated metabolic switch pattern (GLUT1, PGK1, VEGFA), cellular remodeling (BNIP3, HILPDA, P4HA1), and trophic control (IGFBP2/5) in hypoxic niches. |
| 5 | Neuronal signature | ANK3, NAV3, SNHG14, GRIA2, CELF4, FAM155A, FGF14, NLGN1, PPFIA2, DLG2 | This cluster is enriched with canonical neuronal transcripts found in axons (ANK3, NAV3) and synapses (FGF14, GRIA2, DLG2, NLGN1, PPFIA2), possibly indicating synaptophagic astrocytes |

**Supplementary Figure 1.**
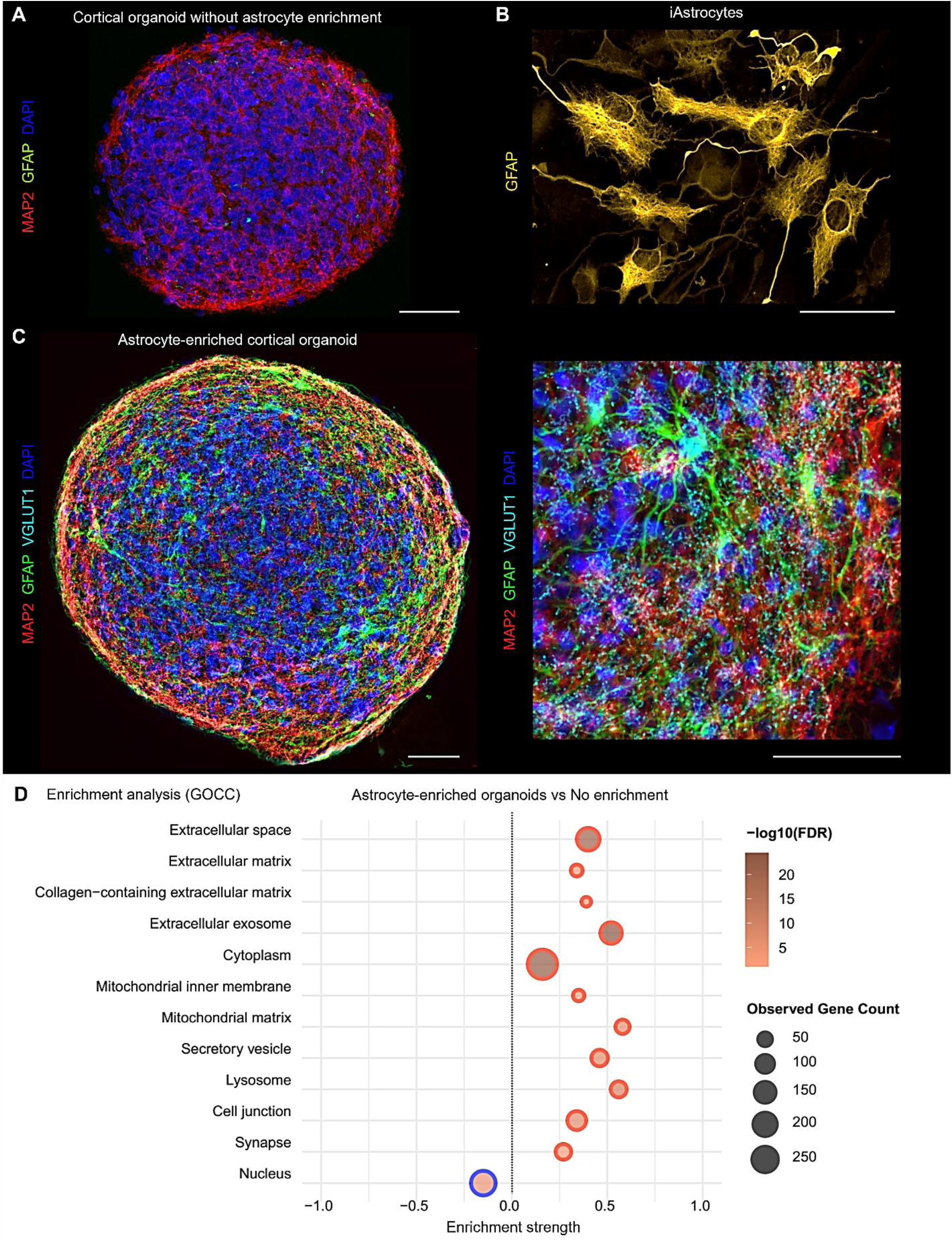
Generation of astrocyte-enriched human cortical organoids. (A) Without astrocyte enrichment, cortical organoids at day 45 contained few astrocytes (GFAP, green). (B) For astrocyte enrichment, pre-differentiated human induced astrocytes (iAstrocytes, GFAP, yellow) were co-cultured with neural precursor cells (NPCs) during organoid formation. (**C**) Astrocyte-enriched organoids develop networks of ramified astocytes (GFAP, green), mature neurons (MAP2, red), and establish glutamatergic synapses (VGLUT1m cyan) at 45 days of cultivation. (**D**) Gene ontology cellular component analysis (GOCC) based on significantly regulated proteins (p < 0.05, fold change difference > 1.5, t-test) measured with LC-MS/MS. The analysis shows upregulated extracellular matrix, secretory programs, and synaptic proteins in astrocyte-enriched vs non-enriched organoids (n=6). Scale bars, 50 µm.

**Supplementary Figure 2.**
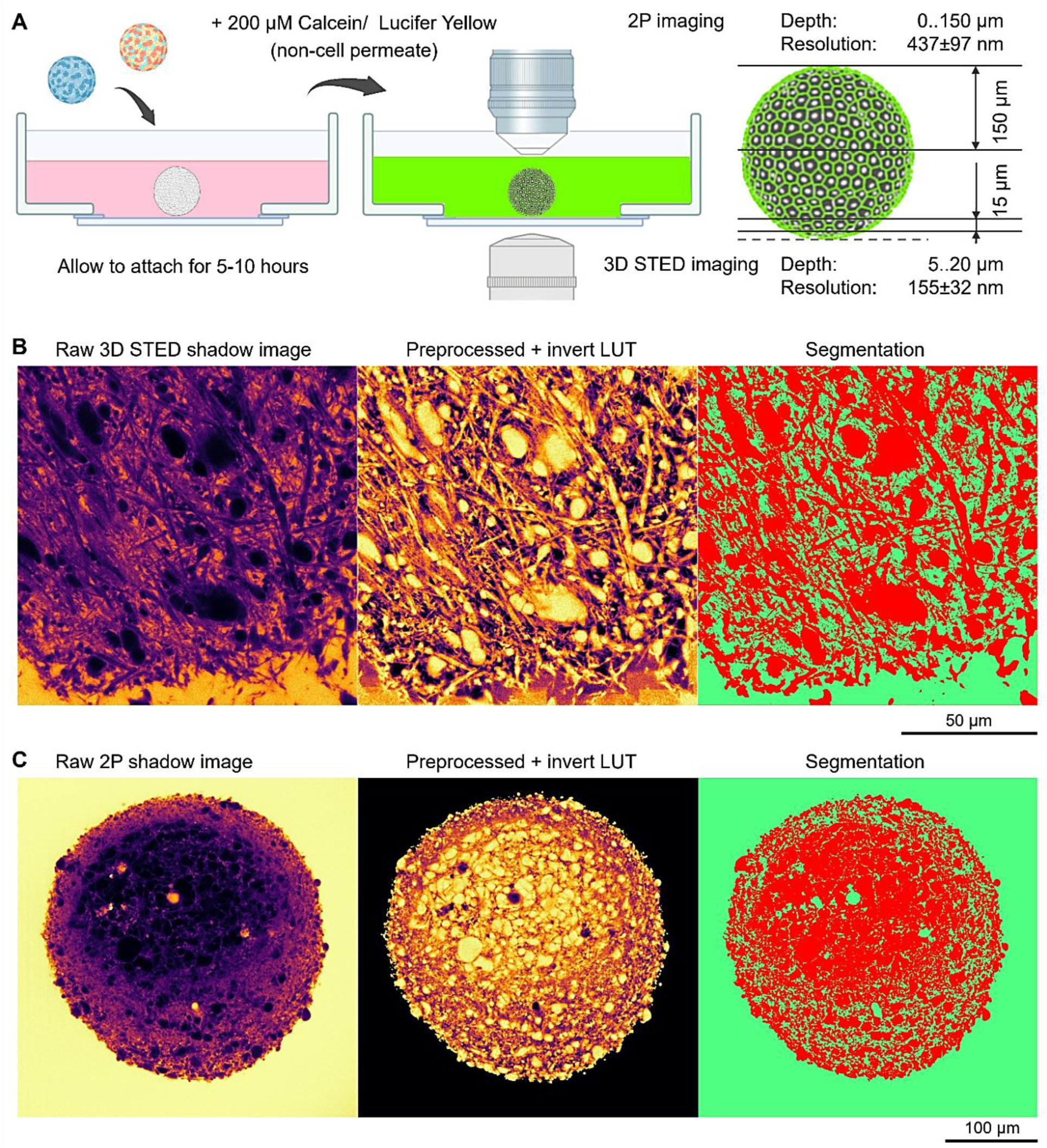
Shadow imaging and image processing. (A) Strategy for visualizing the organoid extracellular space (ECS). Organoids were attached to glass-bottom imaging dishes coated with 1:30 diluted Matrigel for stability. 3D STED microscopy was performed from the bottom of the organoid, starting 5 µm above the attachment site to avoid potential artifacts from Matrigel coating. Apparent resolution at 10 µm depth was estimated by measuring the full width at half-maximum (FWHM) of the smallest visible structures. 2P microscopy was performed from the top of the organoid, and resolution at 100 µm depth was estimated by FWHM of the smallest detectable structures. (**B, C**) Segmentation of cellular (red) and extracellular (green) organoid compartments for ECS volume quantifications in 3D STED and 2P shadow imaging.

**Supplementary Figure 3.**
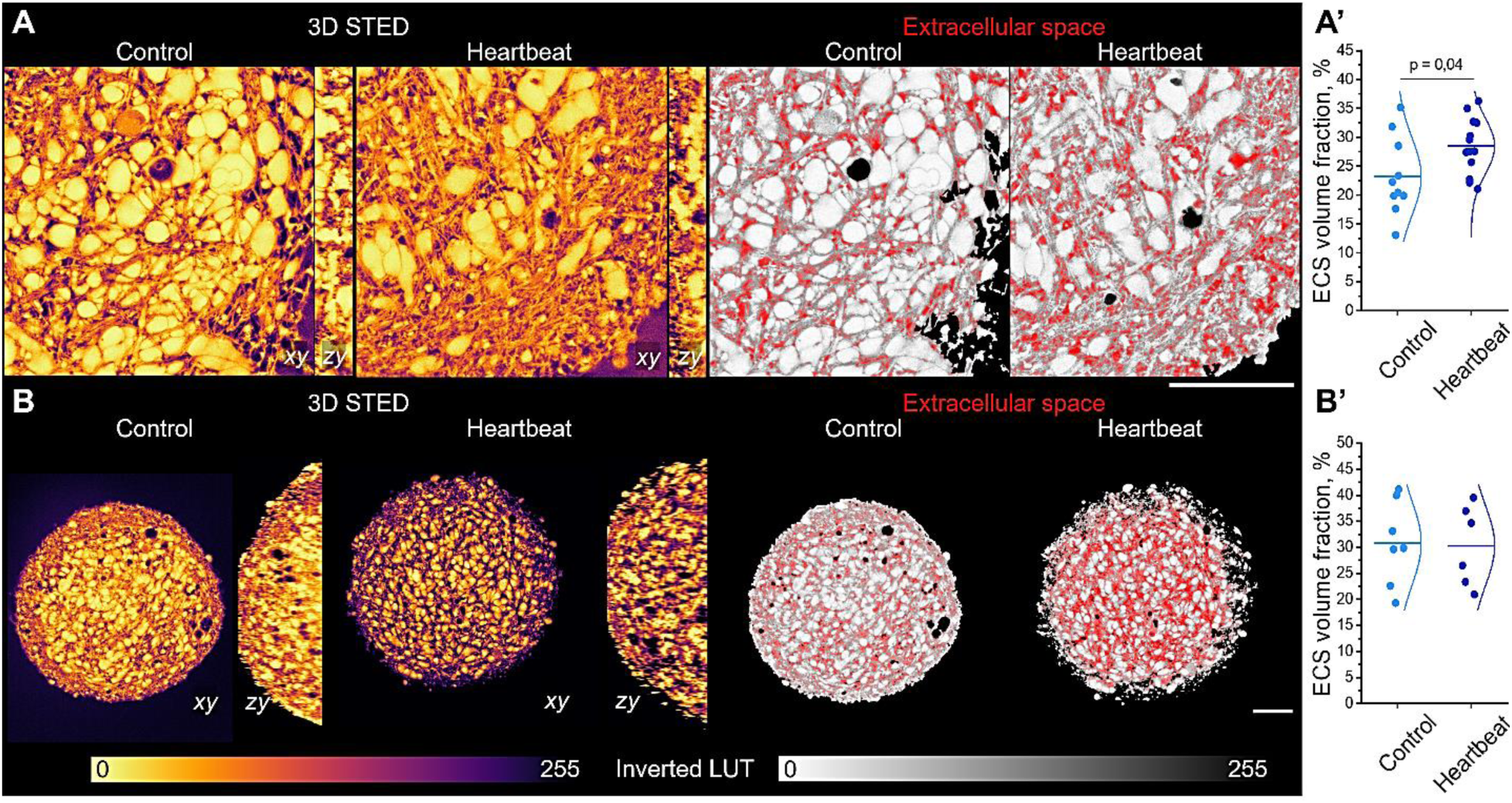
Shadow imaging and ECS volume in human cortical organoids without astrocyte enrichment. **(A)** Superresolution 3D STED shadow imaging and ECS segmentation (red) in live organoids. (**A’**) ECS volume fraction comparison, t-test (n=10-15). (**B**) 2P shadow imaging and ECS segmentation (red) in live organoids. (**B’**) ECS volume fraction comparison, t-test (n=5-6). (**C**, **D**) Images are xy and zy projections; LUT inverted. Scale bars, 50 µm.

**Supplementary Figure 4.**
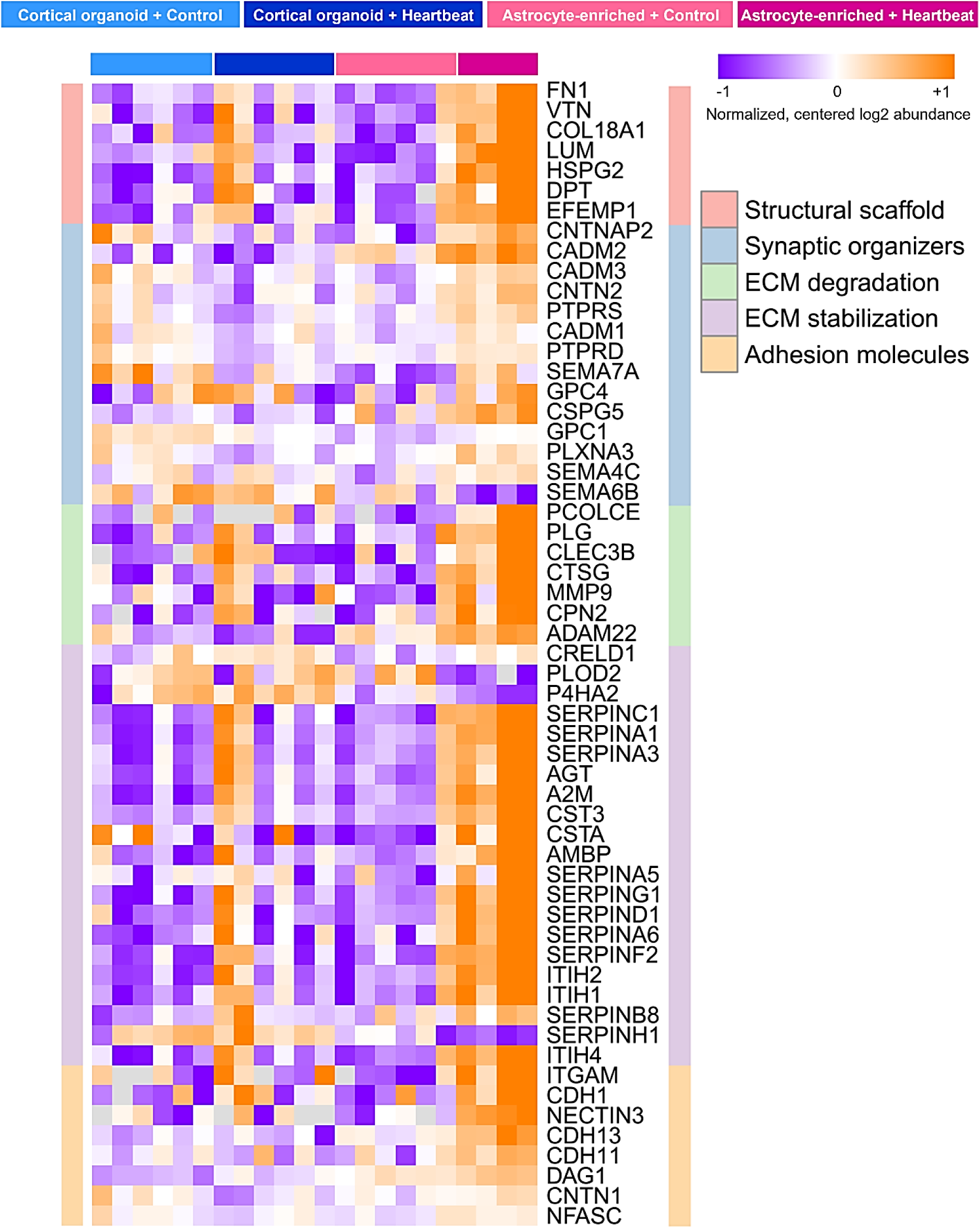
ECM proteins regulated by heartbeat. Heatmap shows normalized, centered log2 protein abundances for ECM proteins found significantly different (p<0.05, t-test) between astrocyte-enriched+control versus astrocyte-enriched+heartbeat groups.

**Supplementary Figure 5.**
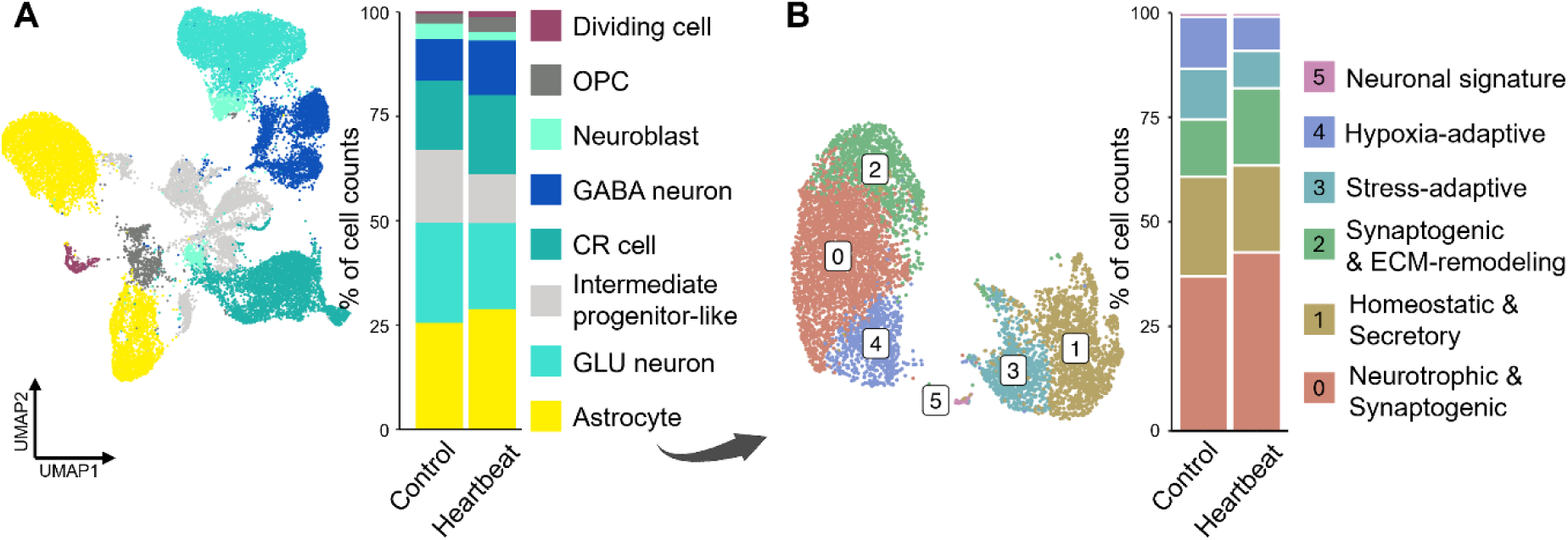
Cellular composition of cortical organoids without astrocyte enrichment. (A) Transcriptomic UMAP map generated from scRNAseq data shows key cellular lineages. (**B**) Astrocyte subtypes revealed by cluster analysis of astrocytic transcriptomes (see **Fig. S6** and **Supplementary Table 2** for functional interpretation).

**Supplementary Figure 6.**
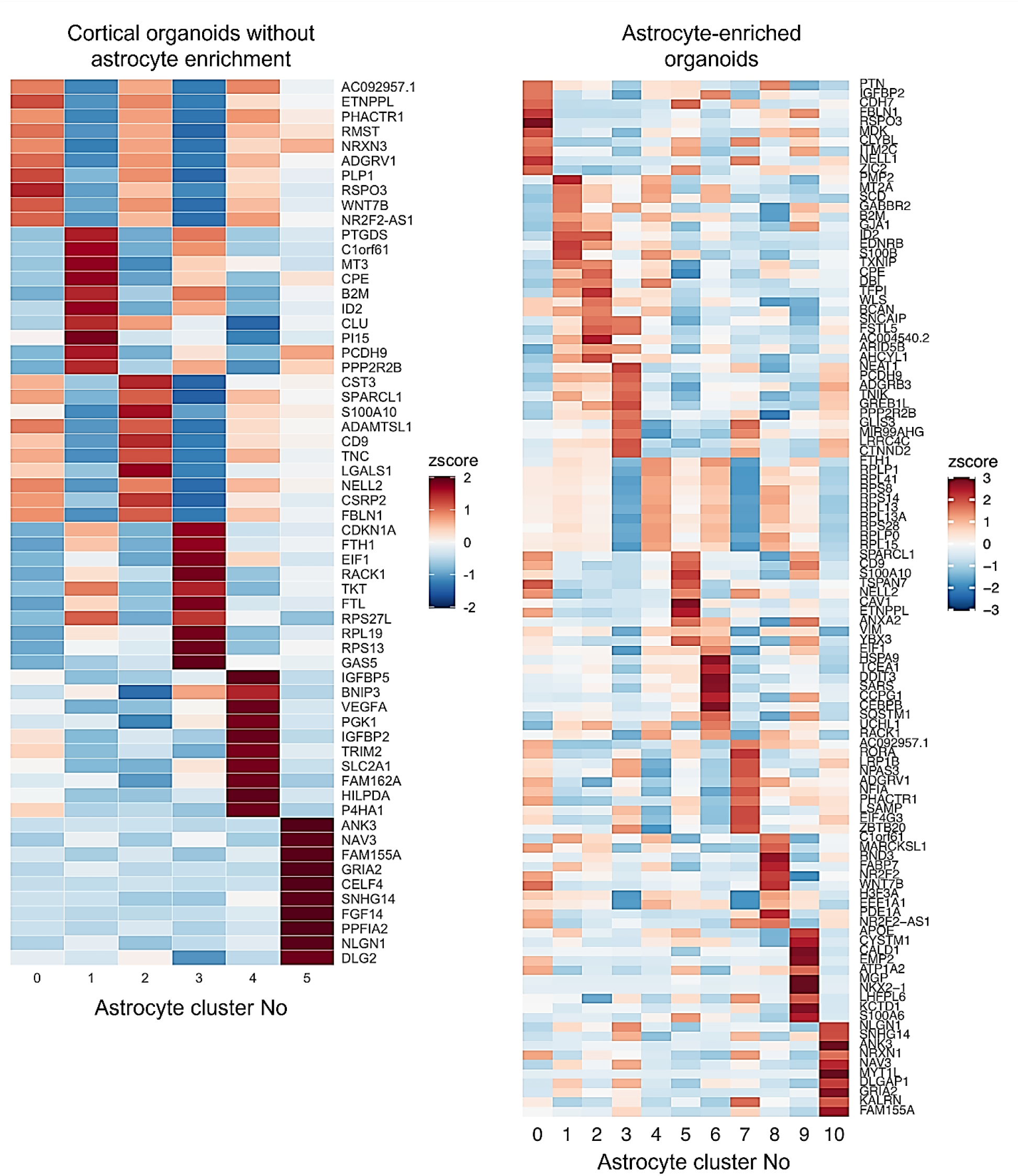
Astrocytic subpopulations identified by scRNA sequencing. Heatmaps show z-scores of key genes differentially expressed in astrocytic subpopulations identified by unsupervised cluster analysis. For functional interpretation, see **Supplementary Table 2**.

**Supplementary Figure 7.**
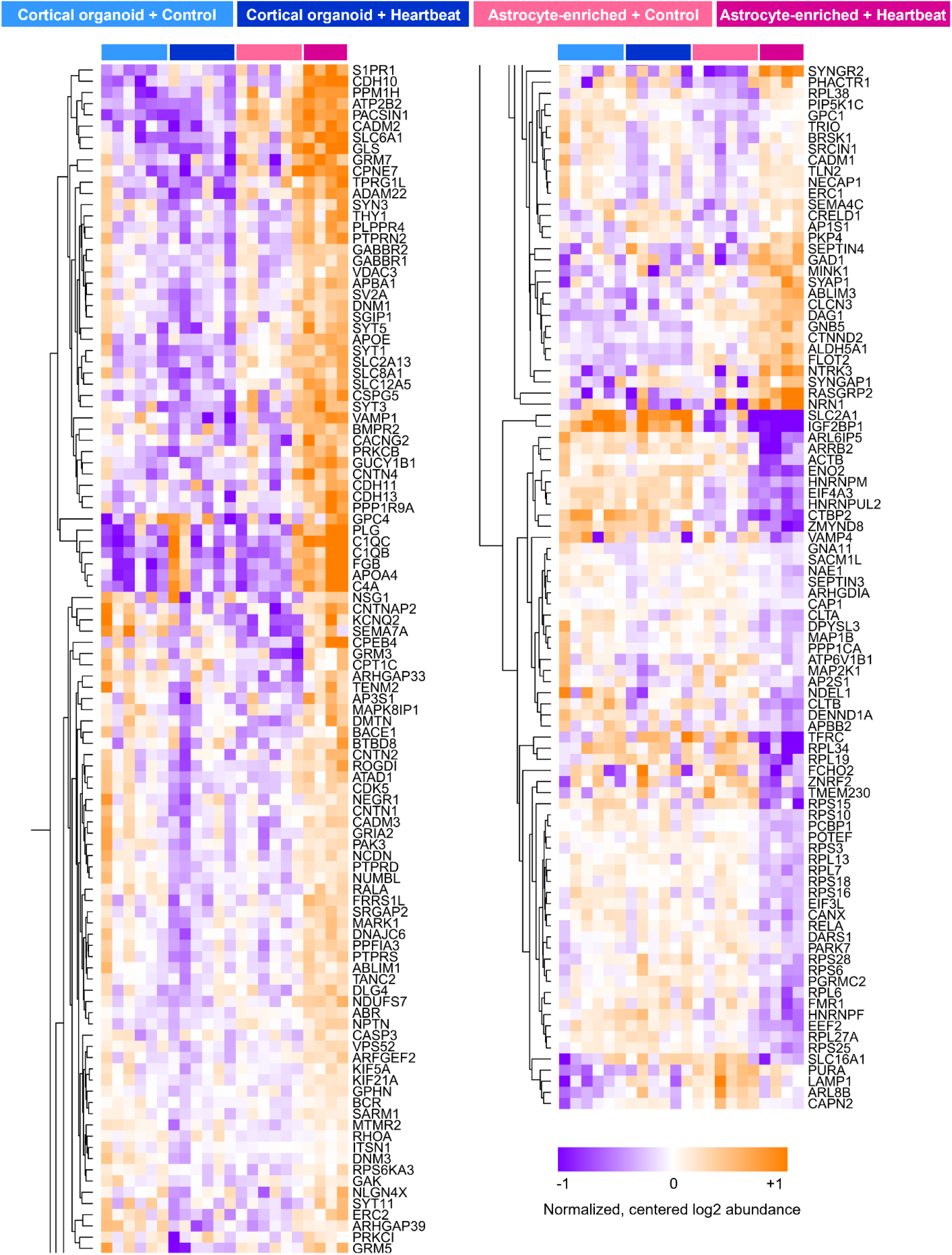
Synapse-associated proteins regulated by heartbeat. Heatmap shows normalized, centered log2 protein abundances for synapse-regulating proteins found significantly different (p<0.05, t-test) between astrocyte-enriched + control versus astrocyte-enriched + heartbeat groups.

**Supplementary Figure 8.**
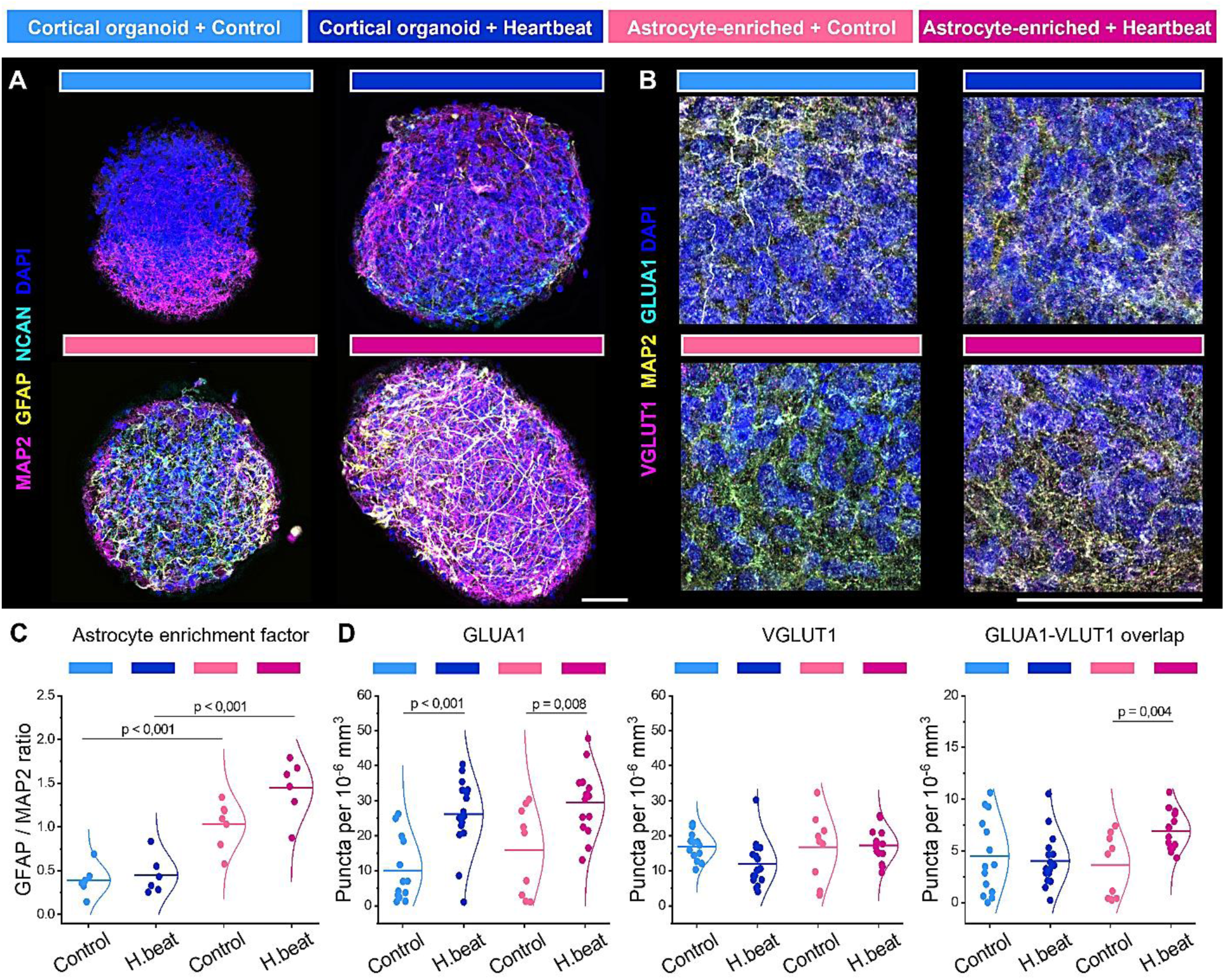
Synaptogenesis in astrocyte-enriched and non-enriched organoids. (A) Immunolabeling of neurons (MAP2, magenta), astrocytes (GFAP, yellow), and neurocan (NCAN, cyan) indicates the enrichment of ramified astrocytes, quantified in (**C**) as the GFAP/MAP2 fluorescence intensity ratio (n=6). (**B**) Immunohistochemical analysis of synaptic markers VGLUT1 (magenta) and AMPA receptor subunit GLUA1 (cyan). Neuronal dendrites are labeled with MAP2 (yellow). (**A, B**) Nuclei (blue) are counterstained with DAPI. Scale bars, 50 µm. (**D**) Quantifications of GLUA1, VGLUT1 synaptic puncta, and their overlaps. (n=9). (**C, D**) Intergroup differences were analyzed by two-way ANOVA followed by t-tests.

**Supplementary Figure 9.**
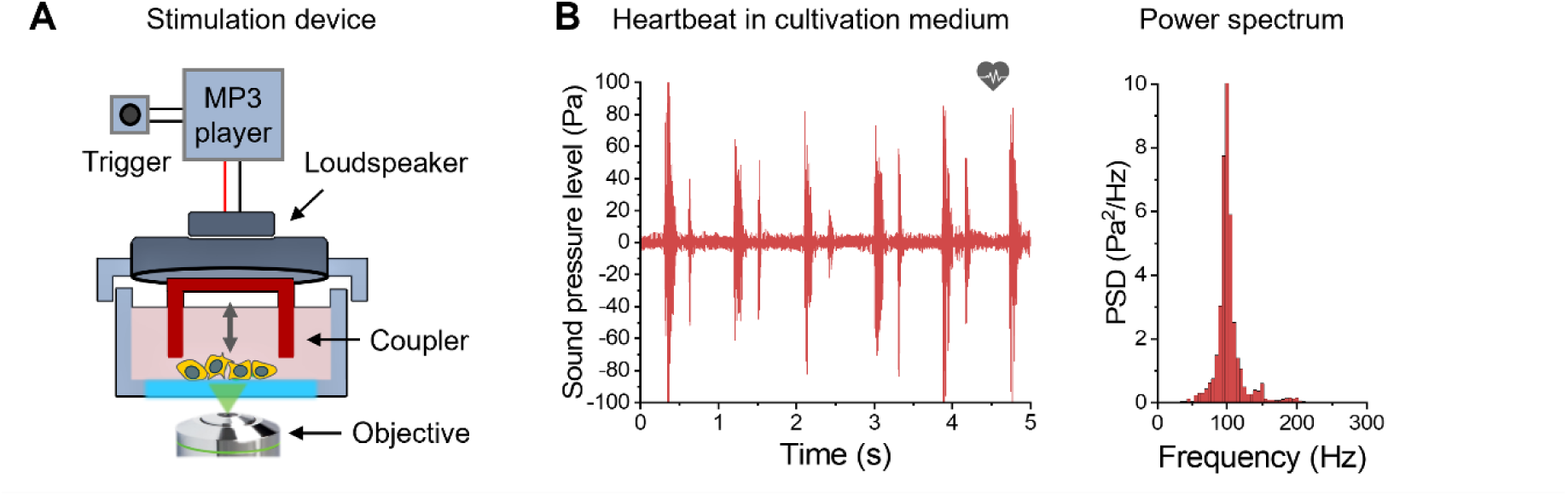
Direct acoustic stimulation in the medium. (A) Custom-made device for direct acoustic stimulation in the medium. A miniature loudspeaker (8 Ohm, 1 Watt) was attached to the lid of an Ibidi dish used for confocal imaging. To achieve the direct transfer of acoustic waves produced by the loudspeaker, a circular coupler was glued to the loudspeaker membrane. The recorded heartbeat was played via an MP3 player with a trigger. (B) Hydrophone recording and power spectrum of the applied heartbeat stimulation.

**Supplementary Figure 10.**
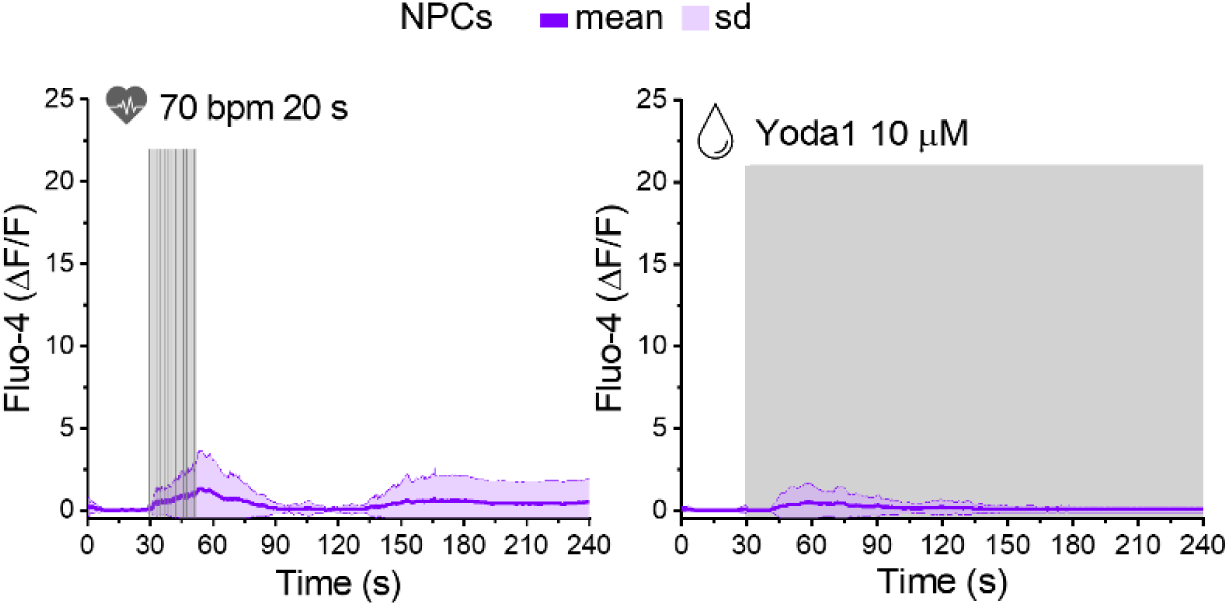
Ca^2+^ responses to heartbeat sound stimulation in NPCs. Averaged Ca^2+^ responses evoked by heartbeat and 10 µM Yoda1 application in NPCs were measured using the Fluo-4 Ca^2+^ indicator. (14-20 cells, n=5)

**Supplementary Figure 11.**
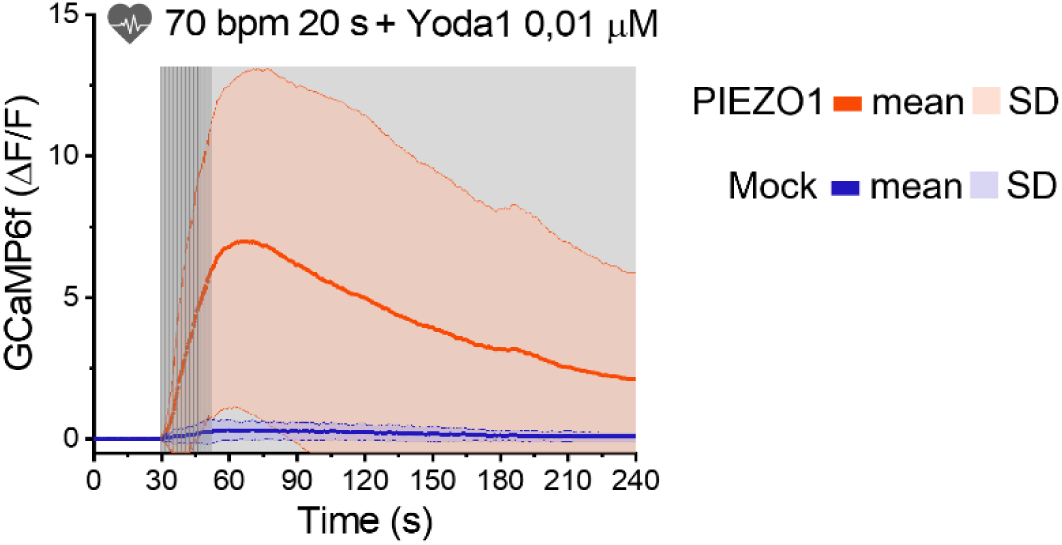
Ca^2+^ responses to heartbeat sound stimulation in HEK293 cells. Averaged Ca^2+^ responses evoked by co-application of 0.01 µM Yoda1 and heartbeat sound stimulation (47 cells, n=7)

**Supplementary Figure 12.**
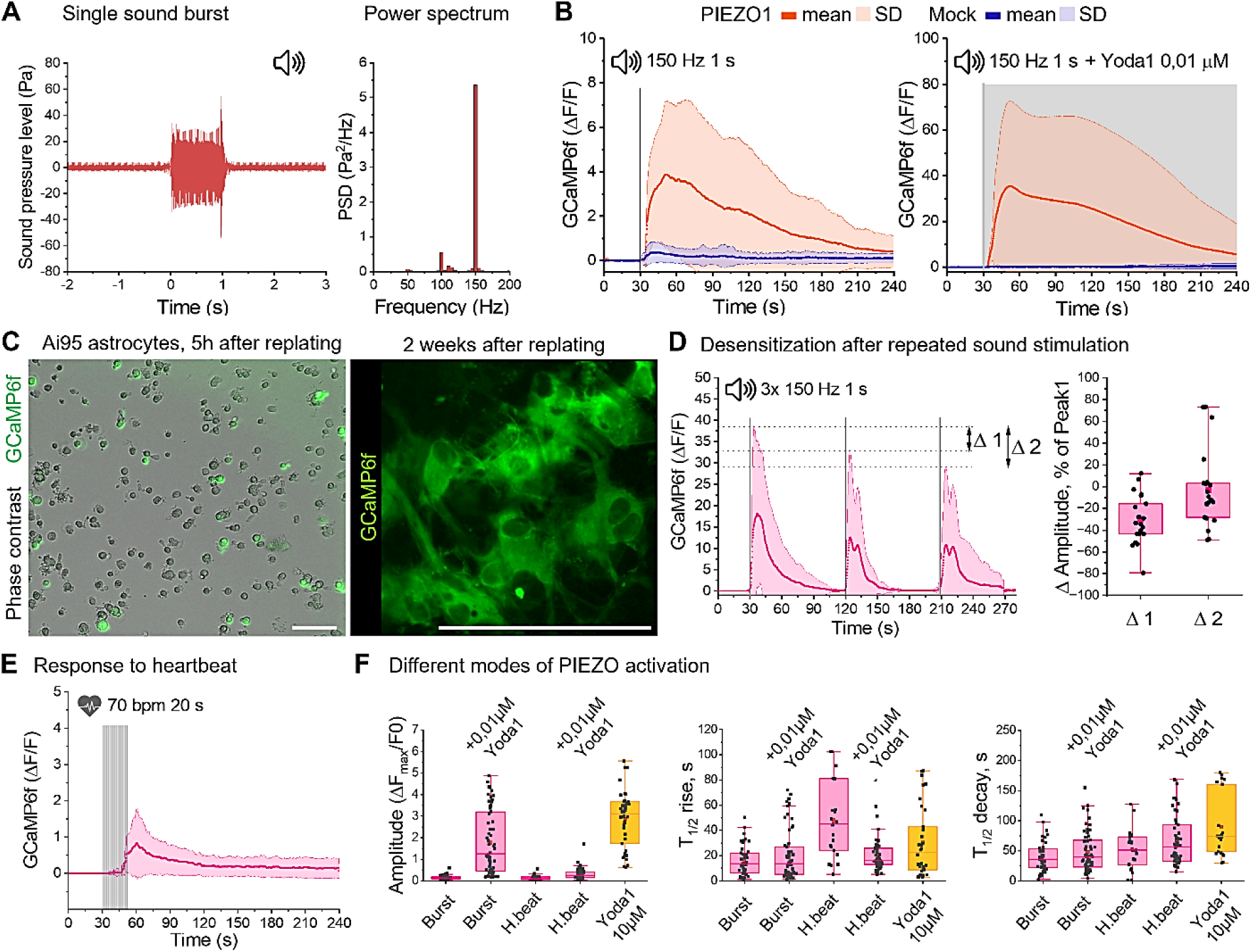
Ca^2+^ responses to low-frequency sound stimulation in primary mouse astrocytes. **(A)** Hydrophone recording and power spectrum of short 150 Hz frequency sound burst stimulation. (**B**) Averaged Ca^2+^ responses to short (1 s) sound bursts in HEK293-P1KO cells co-transfected with PIEZO1-mScarlet (orange) or Mock control (blue) and GCaMP6f. Co-application of a non-activating 0.01 µM Yoda1 concentration and sound bursts amplified PIEZO-mediated Ca^2+^ responses (49-62 cells, n=7). (**C**) Primary astrocyte cultures were prepared from Ai95 (Rosa-CAG-LSL-GCaMP6f) mice, and GCaMP6f expression was induced by TAT-iCre recombinase. One week later, astrocytes were replated onto glass-bottom imaging dishes, and Ca^2+^ imaging was performed in monolayers 2 weeks after replating. Scale bars, 100 µm. (**D**) Ca^2+^ responses to three consecutive 1 s sound bursts delivered at 90-s intervals showed reduced amplitudes compared to the first response, indicating desensitization (21 cells, n=10). (**E**) Averaged Ca^2+^ responses to heartbeat stimulation in primary astrocytes (41 cells, n=12). (**D**) Comparison of different modes of PIEZO1 activation in astrocytes: a single 150-Hz sound burst with (62 cells, n=7) or without (43 cells, n=7) 0.01 µM Yoda1, heartbeat-like stimulation with (54 cells, n=8) or without (23 cells, n=4) 0.01 µM Yoda1, and 10 µM Yoda1 without acoustic stimulation (39 cells, n=5).

**Supplementary Figure 13.**
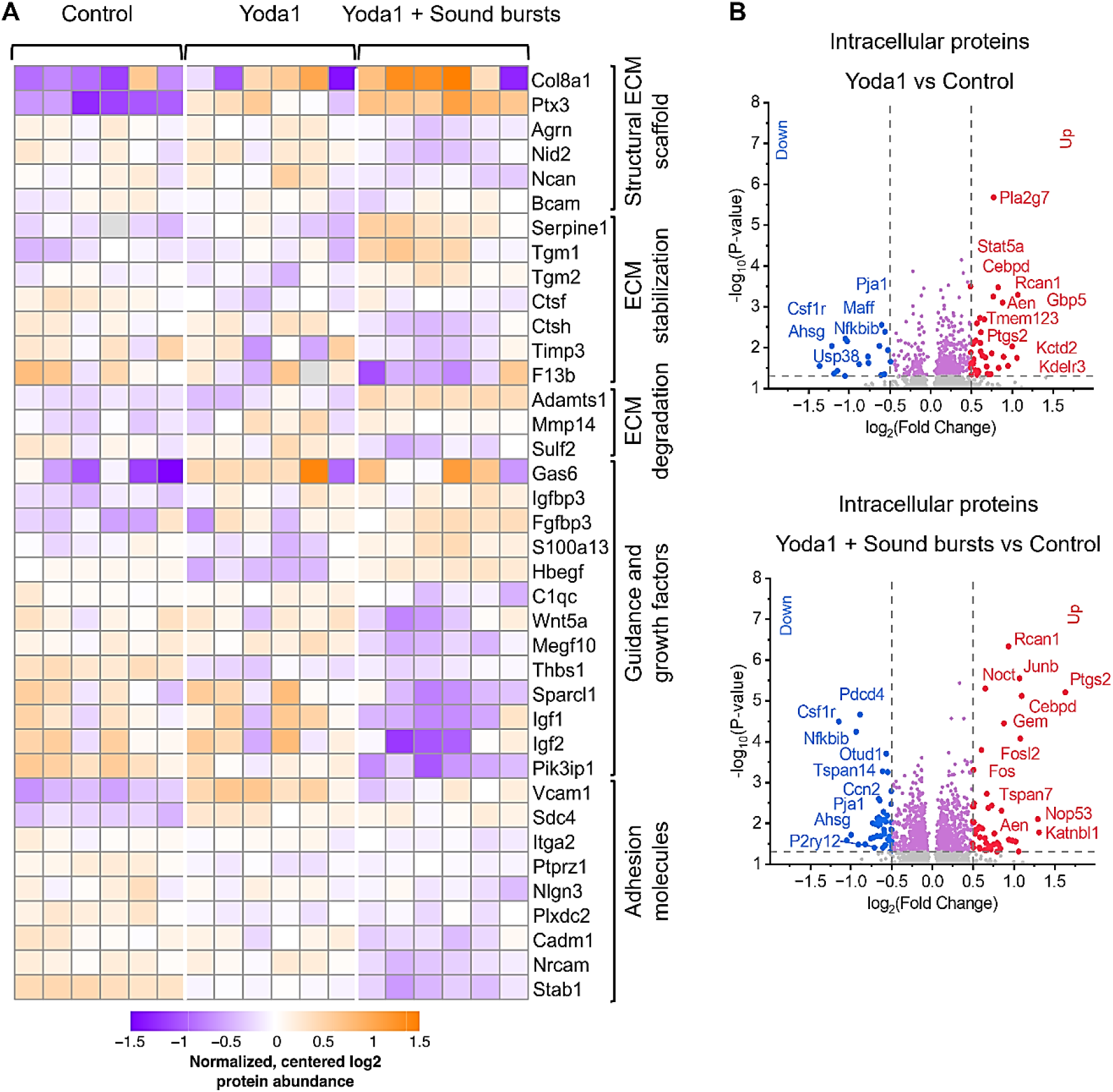
Protein expression changes induced by PIEZO1 activation in primary mouse astrocytes. (A) Heatmap shows normalized, centered log2 protein abundances for ECM proteins found significantly different (p<0.05, t-test) between Yoda1 + sound burst stimulation versus Yoda1 stimulation versus control astrocyte cultures, measured by LC-MS/MS. (**B**) Volcano plots show major regulations of intracellular proteins induced in mouse astrocytes by Yoda1 + sound burst stimulation and Yoda1 stimulation versus control, detected my LC-MS/MS.

## References

Aibar, S., González-Blas, C.B., Moerman, T., Huynh-Thu, V.A., Imrichova, H., Hulselmans, G., Rambow, F., Marine, J.C., Geurts, P., Aerts, J., et al. (2017). SCENIC: single-cell regulatory network inference and clustering. Nat Methods 14, 1083–1086.

Alam El Din, D.-M., Moenkemoeller, L., Loeffler, A., Habibollahi, F., Schenkman, J., Mitra, A., van der Molen, T., Ding, L., Laird, J., Schenke, M., et al. (2025). Human neural organoid microphysiological systems show the building blocks necessary for basic learning and memory. Communications Biology 8, 1237.

Birey, F., Andersen, J., Makinson, C.D., Islam, S., Wei, W., Huber, N., Fan, H.C., Metzler, K.R.C., Panagiotakos, G., Thom, N., et al. (2017). Assembly of functionally integrated human forebrain spheroids. Nature 545, 54–59.

Borbor, M., Yin, D., Brockmeier, U., Wang, C., Doeckel, M., Pillath-Eilers, M., Kaltwasser, B., Hermann, D.M., and Dzyubenko, E. (2023). Neurotoxicity of ischemic astrocytes involves STAT3-mediated metabolic switching and depends on glycogen usage. Glia 71, 1553–1569.

Chi, S., Cui, Y., Wang, H., Jiang, J., Zhang, T., Sun, S., Zhou, Z., Zhong, Y., and Xiao, B. (2022). Astrocytic Piezo1-mediated mechanotransduction determines adult neurogenesis and cognitive functions. Neuron 110, 2984–2999.e2988.

Coste, B., Mathur, J., Schmidt, M., Earley, T.J., Ranade, S., Petrus, M.J., Dubin, A.E., and Patapoutian, A. (2010). Piezo1 and Piezo2 are essential components of distinct mechanically activated cation channels. Science 330, 55–60.

Crapser, J.D., Spangenberg, E.E., Barahona, R.A., Arreola, M.A., Hohsfield, L.A., and Green, K.N. (2020). Microglia facilitate loss of perineuronal nets in the Alzheimer’s disease brain. EBioMedicine 58, 102919.

Dealessandri, G., and Vivalda, M. (2018). The mother’s womb acoustic environment: study of the original sounds and replication for pre-term infants. Journal of Physics: Conference Series 1075, 012056.

Dityatev, A., Schachner, M., and Sonderegger, P. (2010). The dual role of the extracellular matrix in synaptic plasticity and homeostasis. Nat Rev Neurosci 11, 735–746.

Dubin, A.E., Murthy, S., Lewis, A.H., Brosse, L., Cahalan, S.M., Grandl, J., Coste, B., and Patapoutian, A. (2017). Endogenous Piezo1 Can Confound Mechanically Activated Channel Identification and Characterization. Neuron 94, 266–270.e263.

Dzyubenko, E., and Hermann, D.M. (2023). Role of glia and extracellular matrix in controlling neuroplasticity in the central nervous system. Semin Immunopathol 45, 377–387.

Dzyubenko, E., Prazuch, W., Pillath-Eilers, M., Polanska, J., and Hermann, D.M. (2021). Analysing Intercellular Communication in Astrocytic Networks Using “Astral”. Front Cell Neurosci 15, 689268.

Dzyubenko, E., Rozenberg, A., Hermann, D.M., and Faissner, A. (2016). Colocalization of synapse marker proteins evaluated by STED-microscopy reveals patterns of neuronal synapse distribution in vitro. J Neurosci Methods 273, 149–159.

Dzyubenko, E., Willig, K.I., Yin, D., Sardari, M., Tokmak, E., Labus, P., Schmermund, B., and Hermann, D.M. (2023). Structural changes in perineuronal nets and their perforating GABAergic synapses precede motor coordination recovery post stroke. Journal of Biomedical Science 30, 76.

Evans, E.L., Cuthbertson, K., Endesh, N., Rode, B., Blythe, N.M., Hyman, A.J., Hall, S.J., Gaunt, H.J., Ludlow, M.J., Foster, R., et al. (2018). Yoda1 analogue (Dooku1) which antagonizes Yoda1-evoked activation of Piezo1 and aortic relaxation. Br J Pharmacol 175, 1744–1759.

Faissner, A., and Reinhard, J. (2015). The extracellular matrix compartment of neural stem and glial progenitor cells. Glia 63, 1330–1349.

Fawcett, J.W., Fyhn, M., Jendelova, P., Kwok, J.C.F., Ruzicka, J., and Sorg, B.A. (2022). The extracellular matrix and perineuronal nets in memory. Mol Psychiatry 27, 3192–3203.

Gerhardt, K.J., Abrams, R.M., and Oliver, C.C. (1990). Sound environment of the fetal sheep. American Journal of Obstetrics and Gynecology 162, 282–287.

Gottschling, C., Dzyubenko, E., Geissler, M., and Faissner, A. (2016). The Indirect Neuron-astrocyte Coculture Assay: An In Vitro Set-up for the Detailed Investigation of Neuron-glia Interactions. J Vis Exp, e54757.

Hao, Y., Stuart, T., Kowalski, M.H., Choudhary, S., Hoffman, P., Hartman, A., Srivastava, A., Molla, G., Madad, S., Fernandez-Granda, C., et al. (2024). Dictionary learning for integrative, multimodal and scalable single-cell analysis. Nat Biotechnol 42, 293–304.

Hrabetová, S., Hrabe, J., and Nicholson, C. (2003). Dead-space microdomains hinder extracellular diffusion in rat neocortex during ischemia. J Neurosci 23, 8351–8359.

Hua, Y., Weng, L., Zhao, F., and Rambow, F. (2025). SeuratExtend: streamlining single-cell RNA-seq analysis through an integrated and intuitive framework. Gigascience 14.

Hughes, C.S., Moggridge, S., Müller, T., Sorensen, P.H., Morin, G.B., and Krijgsveld, J. (2019). Single-pot, solid-phase-enhanced sample preparation for proteomics experiments. Nature Protocols 14, 68–85.

Jammal Salameh, L., Bitzenhofer, S.H., Hanganu-Opatz, I.L., Dutschmann, M., and Egger, V. (2024). Blood pressure pulsations modulate central neuronal activity via mechanosensitive ion channels. Science 383, eadk8511.

Korsunsky, I., Millard, N., Fan, J., Slowikowski, K., Zhang, F., Wei, K., Baglaenko, Y., Brenner, M., Loh, P.R., and Raychaudhuri, S. (2019). Fast, sensitive and accurate integration of single-cell data with Harmony. Nat Methods 16, 1289–1296.

Kosnoff, J., Yu, K., Liu, C., and He, B. (2024). Transcranial focused ultrasound to V5 enhances human visual motion brain-computer interface by modulating feature-based attention. Nature Communications 15, 4382.

Kumeta, M., Otani, M., Toyoda, M., and Yoshimura, S.H. (2025). Acoustic modulation of mechanosensitive genes and adipocyte differentiation. Communications Biology 8, 595.

Lancaster, M.A., Renner, M., Martin, C.-A., Wenzel, D., Bicknell, L.S., Hurles, M.E., Homfray, T., Penninger, J.M., Jackson, A.P., and Knoblich, J.A. (2013). Cerebral organoids model human brain development and microcephaly. Nature 501, 373–379.

Lewis, A.H., Cui, A.F., McDonald, M.F., and Grandl, J. (2017). Transduction of Repetitive Mechanical Stimuli by Piezo1 and Piezo2 Ion Channels. Cell reports 19, 2572–2585.

Nikolaev, Y.A., Dosen, P.J., Laver, D.R., van Helden, D.F., and Hamill, O.P. (2015). Single mechanically-gated cation channel currents can trigger action potentials in neocortical and hippocampal pyramidal neurons. Brain Res 1608, 1–13.

Onesto, M.M., Kim, J.-i., and Pasca, S.P. (2024). Assembloid models of cell-cell interaction to study tissue and disease biology. Cell Stem Cell 31, 1563–1573.

Paşca, A.M., Sloan, S.A., Clarke, L.E., Tian, Y., Makinson, C.D., Huber, N., Kim, C.H., Park, J.-Y., O’Rourke, N.A., Nguyen, K.D., et al. (2015). Functional cortical neurons and astrocytes from human pluripotent stem cells in 3D culture. Nature Methods 12, 671–678.

Querleu, D., Renard, X., Versyp, F., Paris-Delrue, L., and Crèpin, G. (1988). Fetal hearing. Eur J Obstet Gynecol Reprod Biol 28, 191–212.

Ranade, S.S., Qiu, Z., Woo, S.-H., Hur, S.S., Murthy, S.E., Cahalan, S.M., Xu, J., Mathur, J., Bandell, M., Coste, B., et al. (2014). Piezo1, a mechanically activated ion channel, is required for vascular development in mice. Proceedings of the National Academy of Sciences 111, 10347–10352.

Schneider, C.A., Rasband, W.S., and Eliceiri, K.W. (2012). NIH Image to ImageJ: 25 years of image analysis. Nature Methods 9, 671–675.

Sloan, S.A., Darmanis, S., Huber, N., Khan, T.A., Birey, F., Caneda, C., Reimer, R., Quake, S.R., Barres, B.A., and Paşca, S.P. (2017). Human Astrocyte Maturation Captured in 3D Cerebral Cortical Spheroids Derived from Pluripotent Stem Cells. Neuron 95, 779–790.e776.

Solis, A.G., Bielecki, P., Steach, H.R., Sharma, L., Harman, C.C.D., Yun, S., de Zoete, M.R., Warnock, J.N., To, S.D.F., York, A.G., et al. (2019). Mechanosensation of cyclical force by PIEZO1 is essential for innate immunity. Nature 573, 69–74.

Sykova, E., and Nicholson, C. (2008). Diffusion in brain extracellular space. Physiol Rev 88, 1277–1340.

Tønnesen, J., Hrabĕtová, S., and Soria, F.N. (2023). Local diffusion in the extracellular space of the brain. Neurobiology of Disease 177, 105981.

Tonnesen, J., Inavalli, V., and Nagerl, U.V. (2018). Super-Resolution Imaging of the Extracellular Space in Living Brain Tissue. Cell 172, 1108–1121 e1115.

Tort, A.B.L., Laplagne, D.A., Draguhn, A., and Gonzalez, J. (2025). Global coordination of brain activity by the breathing cycle. Nature Reviews Neuroscience 26, 333–353.

Trujillo, C.A., Gao, R., Negraes, P.D., Gu, J., Buchanan, J., Preissl, S., Wang, A., Wu, W., Haddad, G.G., Chaim, I.A., et al. (2019). Complex Oscillatory Waves Emerging from Cortical Organoids Model Early Human Brain Network Development. Cell Stem Cell 25, 558–569.e557.

Vince, M.A., Armitage, S.E., Baldwin, B.A., Toner, J., and Moore, B.C.J. (1982). The Sound Environment of the Foetal Sheep. Behaviour 81, 296–315.

von Bartheld, C.S., Bahney, J., and Herculano-Houzel, S. (2016). The search for true numbers of neurons and glial cells in the human brain: A review of 150 years of cell counting. J Comp Neurol 524, 3865–3895.

Xu, T., Zhang, Y., Li, D., Lai, C., Wang, S., and Zhang, S. (2024). Mechanosensitive Ion Channels Piezo1 and Piezo2 Mediate Motor Responses In Vivo During Transcranial Focused Ultrasound Stimulation of the Rodent Cerebral Motor Cortex. IEEE Transactions on Biomedical Engineering 71, 2900–2910.

Zeng, W.-Z., Marshall, K.L., Min, S., Daou, I., Chapleau, M.W., Abboud, F.M., Liberles, S.D., and Patapoutian, A. (2018). PIEZOs mediate neuronal sensing of blood pressure and the baroreceptor reflex. Science 362, 464–467.

Zhao, Q., Zhou, H., Chi, S., Wang, Y., Wang, J., Geng, J., Wu, K., Liu, W., Zhang, T., Dong, M.-Q., et al. (2018). Structure and mechanogating mechanism of the Piezo1 channel. Nature 554, 487–492.

